# Adaptation of *Methylobacterium extorquens* to alternating carbon sources identifies the regulator CstR as an intersectional hub of cellular carbon metabolic dynamics and stress response

**DOI:** 10.64898/2026.06.30.735679

**Authors:** Eric L. Bruger, Isabella Ikobe, Chandler N. Hellenbrand, Uly Zigmund, Jannell V. Bazurto

## Abstract

Bacteria frequently face challenges adapting to changing environmental conditions, such as shifting resource utilization, to survive and thrive. Methylotrophs capable of growth on reduced single-carbon compounds are prevalent in the phyllosphere (aerial plant surfaces), where they face continual and predictable shifts in the availability of different plant-produced carbon sources. We examined the ability of the methylotroph *Methylobacterium extorquens* PA1 to adapt to repeated shifts between two different carbon and energy sources: the one-carbon compound methanol and the multi-carbon organic acid succinate, both present in the phyllosphere. Evolved lineages of wild-type cells all increased their capacity for rapid transition between the carbon sources through high frequencies of loss-of-function mutations affecting a previously uncharacterized gene, named *cstR* for carbon source transition regulator, which encodes an orphan single-domain response receiver. Characterization showed that mutant strains were more competitive bidirectionally in the succinate-methanol transition. Though evolved populations of the Δ*efgA* and Δ*ttmR* strains, which are defective in the succinate-to-methanol transition, experienced similar phenotypic improvements in carbon-source transitions, we did not observe *cstR* mutations rise to prominence as extensively or frequently in these lineages.

Transcriptomic work revealed loss-of-function to *cstR* impacted expression of genes involved in motility/chemotaxis, energy metabolism, and stress response, among others, suggesting that it coordinates responses to metabolic cues that are prevalent in certain carbon source and growth phase transitions. Loss of *cstR* function did not compromise exogenous formaldehyde tolerance in the Δ*efgA* and Δ*ttmR* mutants, breaking a previously described tradeoff between these two phenotypes. However, this loss did lead to defects under exposure to certain stressors, including heat, desiccation, oxidative stress agents, and particularly pH stress. Altered levels of NAD+/NADH across conditions, improved growth under acidic pH, and diminished ATP and increased mortality under heightened pH together support a model where CstR is responsible for coordinating cell signaling to manage the balance between growth and maintaining stress resilience.

**Importance:** The ability of heterotrophic bacteria to navigate transitions between different carbon sources is critical to their growth and success. Facultative methylotrophs can grow upon multiple types of carbon sources, including reduced single-carbon compounds, and frequently exist in host-beneficial associations on the leaf surfaces of plants. The exposure to the methylotrophic substrate methanol is known to come in temporal bursts for bacteria residing in the phyllosphere. The utilization of methanol is accompanied by obligatory conversion and exposure to the toxic compound formaldehyde. We sought to investigate the capacity of *Methylobacterium extorquens* to adapt to repetitive transitions between multi-carbon succinate and single-carbon methanol, and in the process identified a novel signaling protein: a single-domain response regulator receiver that mediates cellular growth response to carbon source transitions, at the cost of resistance to certain stressors including oxidative stress and alkaline pH exposure.

## INTRODUCTION

Bacteria often face shifting resource availability that requires them to sense and respond effectively to changes in their natural environments. Heterotrophic bacteria in particular share the challenge (and opportunity) of encountering and breaking down diverse and often complex suites of carbon sources. Among heterotrophs, facultative methylotrophs are organisms that can utilize reduced single-carbon compounds such as methanol, a highly abundant volatile organic compound released from plant leaves and phytoplankton (1, 2) for carbon and energy but can also metabolize multi-carbon compounds such as sugars and organic acids (3, 4). The phyllosphere, comprising the aerial surfaces of plants, is a nutrient-poor, rapidly shifting physiochemical environment dominated by bacteria. Here, there are noted temporal fluctuations in carbon availability on plant leaves, due to plant host physiology that includes well-established diurnal pulses of volatile organic compound release, including methanol, controlled by the timed opening and closing of stomata (1, 5–9). In this environment, bacteria experience heightened exposure to ultraviolet radiation, water limitation, temperature variation, and oxidative stress through exposure to ozone, photo-oxidation, quinolone antibiotics, phthalic acid esters, and heavy metals (10–13). Thus, inhabitants of the phyllosphere, such as *Methylobacterium* species, are often metabolically versatile and adept at mitigating environmental stressors (10, 14).

Signaling transduction pathways allow bacteria to maintain a timely and coordinated response to environmental cues, transmitting an environmental signal into a functional response. In their simplest form, a single protein has a sensory input domain that senses the signal and an output domain (often DNA-binding) that alters gene expression accordingly. In a two-component system, these functions reside in two distinct proteins, and the signal is transduced from a sensory protein to a response regulator protein via phosphorelay, while multi-component systems are more complex, branched, and involve intermediate proteins that can integrate multiple input signals. *Methylobacterium extorquens*, a beneficial plant-associated bacterium that promotes plant health, encodes numerous sensory proteins (213 total)(15). This includes 76 annotated/predicted response regulators, 29 of which are single-domain proteins (single-domain response regulators, “SD-RRs”)(16–18) that lack an effector output domain. Though many of these predicted SD-RR genes exist in the *M. extorquens* genome, few have been assigned specific physiological roles. The SD-RR encoded by *Mext_0407* was previously noted to be involved in the cell’s general stress response, which is a common function SD-RRs have been reported to perform (19). Though the full biological significance and specific cellular and behavioral functionality provided by these SD-RRs is still being uncovered, these proteins have been demonstrated to impact diverse traits that include chemotaxis, motility, spatial coordination, stress response, and allosteric regulation in other well-studied bacterial systems to provide effective sensing and response to complex, multi-faceted natural environments (18).

*M. extorquens* experiences growth delays upon the transition from multiple-carbon heterotrophic to single-carbon methylotrophic growth, compared to carbon transitions in the opposing direction or to continued growth on one carbon source, thought to be mediated by endogenous formaldehyde production (20, 21). The growth delay is exacerbated in strains that are lacking the formaldehyde sensor EfgA and the formaldehyde-responsive TtmR transcription factor. Both strains are differently defective in metabolic homeostasis in comparison to wild-type (WT), with the Δ*efgA* strain experiencing inflated formaldehyde pools and the Δ*ttmR* mutant having decreased cellular formate and secreting excess formaldehyde. Given this initial offset, we sought to understand whether genotypes with distinctly different phenotypes might improve their respective transitions to methylotrophy upon exposure to frequent and repetitive carbon substrate switching. Can these genotypes reach similar or identical fitness peaks, despite very different starting positions? Would fitness improvements occur via identical or diverse genetic changes? Additionally, given that the loss of *efgA* and *ttmR* confers increased resistance to exogenous formaldehyde, would any such improvements in these genotypes inevitably come at the cost of resistance to exogenous formaldehyde?

To examine the pressures and potential adaptation achieved by cells to shifting carbon resources, to and from methylotrophy, as they may in the phyllosphere environment, we imposed selective evolutionary regimes to cells undergoing repeated alternations of sole carbon sources. Herein, we describe the resulting adaptation to repetitive carbon transitions and identify a novel SD-RR, we rename CstR (**<u>c</u>**arbon **<u>s</u>**ource **t**ransition **<u>r</u>**egulator) which was impacted by mutation in all WT lineages, and whose evolved mutations universally swept those populations. We further compare the impact of mutating *cstR* in Δ*efgA* and Δ*ttmR* strains. A straightforward hypothesis would be that distinct mutations would be necessary to restore growth to Δ*efgA* and Δ*ttmR* mutants due to the distinct root causes of their initial physiological defects. Surprisingly, we find that the loss of *cstR* restores their optimal transitions to methylotrophy, even surpassing WT growth. Further, physiological analyses in a WT background identified a tradeoff that emerges when *cstR* function is removed, rendering the resulting cells susceptible to impaired growth and decreased survival under certain abiotic stress conditions, and suggesting that cells have altered internal pH. Ultimately, we saw that Δ*cstR* exhibited improved growth parameters in conditions transitioning between succinate and methanol compared to WT, while experiencing growth deficits under several tested stress conditions, including heat shock, desiccation, hydrogen peroxide, and alkaline pH. Here, we propose a model whereby the altered regulation provided by loss of CstR function leads to cells biased towards growth at the expense of stress tolerance, largely mediated by shifts in fundamental cell properties including energy production, redox balance, and proton homeostasis, ultimately impacting cell viability under strained conditions.

## MATERIALS AND METHODS

### Bacterial strains, media, and chemicals

All strains used in the present study were derived from the wild-type *Methylobacterium extorquens* PA1 strain (22) with *bcsABZC* (*Mext_1367*–*Mext_1370*, yielding the strain CM2730 deleted to eliminate aggregation during growth in liquid culture growth (23), and are listed in **Table S1**. Strains were cultivated using *Methylobacterium* PIPES (piparazine-N,N’-bis(2-ethanesulfonic acid), Santa Cruz Biotech) (‘MPIPES’ or ‘MP’) media supplemented with 3.5 mM succinate (C₄H₄Na₂O₄•6H₂O, CAS# 6106-21-4, Alpha Aesar), 15 mM methanol (CH_4_O, CAS# 67-56-1, Sigma-Aldrich), 20 mM formate (CHO_2_Na, CAS# 141-53-7, Sigma-Aldrich), or 20 mM oxalate (Na_2_C_2_O_4_, CAS# 62-76-0, Sigma-Aldrich) as carbon sources, unless otherwise noted. Solid MP media contained Bacto agar at 15 g/L (Apex Bioresearch Products), and carbon sources succinate at 15 mM, methanol at 125 mM, or oxalate at 50 mM when used. When required for selection in the media, tetracycline was added to a final concentration of 12.5 μg/mL. Where used to induce gene expression from plasmids, 50 μM cumate (4-Isopropylbenzoic acid, CAS Number 536-66-3, Sigma catalog number 268402) was used. **Experimental evolution.** From individual colonies grown on plates of MP media containing succinate, six independent cultures of CM2730 (WT), CM3745 (Δ*efgA*), and CM4732 (Δ*ttmR*) were grown in 5 mL liquid MP medium in batch culture in 18 x 150 mm glass tubes supplemented with 3.5 mM succinate. Upon reaching stationary phase, the first transfer was performed by inoculating 78 μL of culture (1/64 dilution) into fresh MP medium containing 15 mM methanol. Thereafter, cultures were transferred between these two carbon sources in an alternating fashion for 174 generations (29 transfers every 48-72 hours, at 6 generations per transfer, see **Fig. S1** for overall experimental scheme). Aliquots of the populations were harvested at regular intervals (every 2 transfers, or 12 generations) for storage at -80 °C in a suspension containing the cryoprotectant DMSO at 8% final concentration.

### Genomic sequencing

At generation 174 of the succinate-to-methanol evolution experiment, cells were pelleted and genomic DNA was extracted from WT populations 1-3. The samples were sequenced via Illumina sequencing (1x100 bp reads for populations 1 and 2, 2x150 bp reads for population 3). Sequencing reads were processed with FastQC and Trimmomatic (version 0.39)(24, 25) and genomic changes were determined using breseq (26) compared to the annotated reference genome, GenBank accession number CP000908.1. Following identification of parallel mutations to *Mext_1965* (*cstR*), genomic DNA was isolated from all populations and amplicons spanning the *Mext_1965* coding sequence (CDS) and surrounding region (596 nucleotides) were amplified by polymerase chain reaction (PCR) with primers 5’-gttagggctgggaaccggc and 5’-cgcgatcgcgaacggg and submitted for Sanger sequencing (Plasmidsaurus Inc.) to quantify read counts and determine relative allele frequencies. Combined mutant *Mext_1965* allele identities and frequencies are listed in **Table 1**.

**Table 1.**
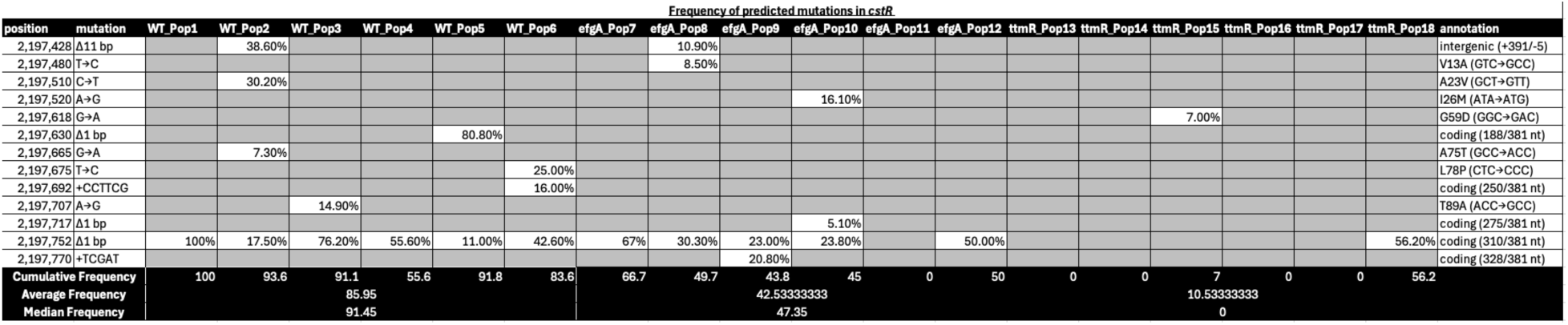
Frequency of *cstR* mutations across experimental populations.

### Genetic approaches

Markerless gene deletions were generated by allelic exchange as previously described with modifications (21, 27). Bi-parental conjugal matings were performed by mixing *Escherichia coli* S17-1 (28) cells carrying the pPS04-based donor plasmid (29) with the pertinent *M. extorquens* recipient strain (both pelleted and resuspended in nutrient broth) in a 1:5 ratio. The mixture was plated on nutrient agar and grown overnight at 30 °C, removed and resuspended in MP medium lacking carbon and nitrogen, serially diluted in identical media, and plated on selective medium supplemented with 15 mM succinate, 5 mM methylamine (non-permissive nitrogen source for *E. coli*), and 50 μg/mL kanamycin. Sucrose counterselection was accomplished by restreaking isolated colonies from the selection plates on MP medium supplemented with 15 mM succinate and 5% sucrose. Successful deletions of the *Mext_1965* locus were identified by PCR screening. Donor plasmids and primers were designed using SnapGene software. Plasmids were assembled using New England Biolabs (NEB) HiFi assembly kits.

### Growth quantification

Isolated colonies were picked from solid agar plates and placed into liquid media. All growth experiments were conducted with at minimum three biological replicates, each individual colony representing a separate biological replicate. Initial cultures were grown in 2 mL liquid MP media supplemented with a sole carbon source. These cultures underwent incubation at 30 ℃ with shaking at 250 rpm and were subsequently subcultured (1/64) into 5 mL of MP medium containing the respective carbon source for acclimation. Following acclimation periods of approximately 48 hr, 10 μL of stationary phase cultures were subcultured (1/64) into 630 μL of MP medium supplemented with the appropriate carbon sources. Unless otherwise stated, this step took place in 48-well polystyrene plates (Falcon, Ref No. 351178), sealed with 18-gauge needle-punctured adhesive optical film (VWR, Cat No. 60941-064) to minimize liquid evaporation and well-to-well variation in aeration. The cultures were then incubated without lids at 30 ℃ with a 2 ℃ gradient to prevent film condensation in a BioTek Epoch 2 microplate spectrophotometer, utilizing single orbital shaking at 800 rpm. Cell density was monitored by measuring optical density at 600 nm (OD_600_) every hour. The cell growth rate (μ) and generation time (*gt*) were calculated through non-linear regression of data points identified within the exponential growth phase, where μ = ln(X_T_/X_0_)/T, and generation time *gt* = ln(2)/μ, with X representing OD_600_, and T denoting elapsed time. Yields were reported as maximum OD_600_ or CFU/mL reached. Lag times were determined through non-linear regression of groups of three consecutive growth readings: statistical non-significance from the non-linear regression of exponential phase cells marked the end of lag phase.

### Competition Assays

Competition experiments were performed as described previously (30) through co-culture of strains in a pairwise fashion, where in one strain was genomically tagged with the fluorescent protein mCherry to differentiate from the other strain. Cultures were combined in equal volumetric ratios with the *hpt*::*mCherry* strain CM3839 (disruption of the gene that encodes the enzyme hypoxanthine-guanine phosphoribosyltransferase). Subsequently, cell suspensions were introduced into 5 mL of MP medium containing a sole carbon source. The cultures were incubated at 30 °C with shaking. Upon reaching stationary phase (48-hour incubation period), 1 mL of each culture was mixed with DMSO (final concentration of 8%) and stored at -80 °C until analysis.

For analysis, the cultures were thawed, subjected to centrifugation, and then resuspended in 1 mL of PBS, followed by a 1:10 dilution before examination of population composition using flow cytometry. The frequency of competitors was assessed by passing mixed population samples from the experiment’s start (F_0_) and end (F_1_) through an LSRFortessa X-20 flow cytometer (Becton Dickinson Biosciences). Approximately 100,000 cells per sample were gated based on forward and side scatter. Competitors were distinguished by comparing fluorescence, excited at 561 nm and emitted through a 610 nm with 20 nm BP filter. Malthusian relative fitness measurements (*w*) were calculated using the methods previously detailed (30).

### Protein structure/function predictions

The sequence of the annotated Mext_1965 protein was aligned with a selection of related sequences from other bacteria including proteins with implicated relevance or belonging to an array of environmental bacteria including from those known to be at high abundance in the phyllosphere. Multiple sequence alignments were conducted via Clustal Omega 0.2.4 (31). AlphaFold Server was used to generate structural predictions (32). Structural comparisons that identified similarity to Rrf1 homolog structures arose from comparisons through the Gaia database (33).

### RNA-sequencing analysis

The wild-type (CM2730) and the Δ*Mext_1965* mutant (JB457) strains were grown in biological triplicate in MP medium with succinate or a succinate-to-methanol transition, as described above (100 mL culture, OD_600_ = 0.25). Cells were harvested by centrifugation in the Eppendorf centrifuge 5810R (5 min, 4,000 x *g*, and 4 °C) and washed with ice-cold MP media (lacking carbon). Pellets were immediately frozen by submerging tubes in liquid nitrogen and stored at -80 °C. Nucleic acid extraction and molecular manipulation (RNA extraction, cDNA generation, and library preparation) and sequencing were conducted by Genewiz (South Plainfield, NJ). Total RNA was extracted with the RNeasy Plus minikit (Qiagen); rRNA was depleted with the Ribo-Zero rRNA removal kit (Illumina); the quality of the resulting RNA samples was determined with an Agilent 2100 bioanalyzer and Qubit assay. Once generated, the cDNA library was sequenced on an Illumina NovaSeq instrument (2x150 bp). The raw data (FASTQ format) was submitted to further analysis. An analysis pipeline developed by the University of Minnesota Genomics Center and the Research Informatics Solutions (RIS) group at the University of Minnesota Supercomputing Institute was used for data analysis (34). The pipeline uses FastQC (25) to assess the quality of the sequencing data, Trimmomatic (24) to remove the low-quality bases and adapter sequences, HISAT2 (35) to align the curated reads to the *M. extorquens* PA1 genome (GenBank accession CP000908.1), Cuffquant and Cufnorm from the Cufflinks package (36, 37) to generate fragments per kilobase per million (FPKM) expression values, and featureCounts from the Rsubread R package (38) to generate raw read counts. DESeq2 was used to convert raw count data to normalized counts, and all further statistical analyses were carried out in R (39). Differentially expressed genes in the Δ*cstR* mutant were identified by a pairwise comparison to the WT strain and had a false discovery rate (FDR) of 0.01. Enrichment among differentially expressed genes (DEGs) was assessed with the ShinyGO 0.77 interface (40).

### Motility assays

Cell cultures of CM2730 and JB457 were prepared as previously mentioned in MP liquid media containing 3.5 mM succinate in biological triplicate. Following acclimation, 10 μL of each culture was ejected into the center of an MP motility plate containing 3.5 mM succinate and 0.25% agar. Plates were incubated at 30 °C and allowed to grow for several days until expansion ceased. Following growth, plates were imaged using an Epson V600 scanner and relative colony areas were determined against a standard reference area and compared using the ImageJ software suite (41).

### Stress tolerance experiments

A variety of potential stressors were applied to *M. extorquens*; the cellular response was assayed by monitoring growth (OD_600_) or survival (CFU/mL) as described above.

#### Ethanol, Salt, Peroxide, Formaldehyde

Cells were exposed to ethanol (up to 6% (v/v)), salt (up to 500 mM), hydrogen peroxide (0.1% (v/v)), and formaldehyde stress (up to 14 mM) during growth on succinate, methanol, or the transition to methylotrophy as indicated. Growth experiments were executed as described above unless otherwise stated. For formaldehyde MIC data, growth was monitored for up to 4 days via OD_600_ to determine ultimate growth.

#### Ultraviolet Radiation

Cells grown to stationary phase were diluted in carbon-less MP media and plate on MP plates with succinate as the carbon source. Once dried, the plates were exposed to ultraviolet light at an intensity of 240 mW/cm^2^ at 1m, with a peak UV Output at 253.7 nm in a sterile biosafety cabinet (SterilGARD III Advance, The Baker Company) for a range of time intervals. The plates were then incubated at 30 °C and enumerated after visible colony formation to determine survival.

#### Desiccation

Cells grown to stationary phase were diluted in carbon-less MP media and plated on mixed cellulose ester filters (Immobilon-NC, MilliporeSigma Catalog Number HATF08550), allowed to dry, and maintained in a dry and dark space for extended intervals of time ranging from 0 days to 8 months. After incubation for the allotted desiccation period, filters were transferred and adhered to MP petri plates containing succinate as the sole carbon source. Plates were incubated at 30 °C and viable cell counts were enumerated to determine survival.

#### Temperature stress

Cells grown to stationary phase were diluted in carbon-less MP media and exposed to heightened temperature in a 50 °C water bath for a range of time intervals. The cultures were then dilution plated onto MP-succinate media, incubated at 30 °C, and enumerated following visible colony formation to determine survival.

#### Stationary phase viability

Cultures of cells growth in MP media containing oxalate as the sole carbon source were inoculated and passaged at 30 °C with shaking and periodically subsampled, serially diluted ten-fold and spot-plated on MP media containing 15 mM succinate, from which viable cell counts were enumerated.

#### pH stress

In the initial oxalate growth experiments, pH was monitored by periodically taking a portion of culture, pelleting out cells, and measuring pH of the remaining supernatant. For pH altered experiments, cells were grown in MP media containing succinate adjusted to a range of final pH values from 4-11. Death curves were evaluated by growing cultures to stationary phase, collecting cells via centrifugation and resuspended in the same volume of MP media lacking carbon with pH adjusted and incubated at 30 °C. Dilution plating was then carried out over time to track survival.

### Metabolite analysis

CM2730 and JB457 were acclimated to stationary phase in MP media containing 3.5 mM succinate and then diluted into 5 mL media in 18x150 mm tubes containing either 1) 3.5 mM succinate, 2) 15 mM methanol, or 3) 20 mM oxalate in biological quadruplicate and incubated at 30 ℃ with shaking at 250 rpm. OD_600_ was measured over time using a Spec20 spectrophotometer to measure growth. Exponential phase samples were harvested at OD_600_ = 0.2 for succinate and methanol cultures, and OD_600_ = 0.1 for oxalate cultures. Stationary phase samples were harvested after 24 hrs of growth for succinate cultures and 40 hours for methanol and oxalate cultures. At sampling time, viability plating was conducted to calculate cell densities. For each sampling point, two cultures were separately centrifuged at 4 ℃, 4200 rpm for 10 minutes. Supernatant was discarded and cell pellets were stored at - 80 ℃. To assess ATP, cells were processed with the Bac-Titer Glo assay kit (Promega) and incubated for 30 minutes in black-walled 96-well plates, after which luminescence was measured in a SpectraMax iX3 plate reader. To assess redox pools, pellets were resuspended and divided in half; one resulting sample was treated with 0.2 M HCl to assess NAD+ concentrations, while the other was treated with 0.2 M NaOH to assess NADH concentrations. After incubating at 60 ℃ for 10 minutes, samples were neutralized, 50 μL were exposed to 50 μL NADH-Glo reagent (Promega) and incubated for 60 minutes in black-walled 96-well plates, followed by luminescence measurement.

### Statistical Analysis

Where appropriate, statistical comparisons were made via two-way ANOVA with Tukey’s HSD posthoc test to account for multiple comparisons. RNA-seq statistical comparisons were carried out as described in that methods section. All experiments are repeated to confirm consistency, and within experiments contain a minimum biological N=3 of samples per strain per condition.

## RESULTS

### Repeated switching to and from methylotrophic growth leads to rapid phenotypic improvements for evolved populations

To examine the capacity for adaptation during carbon transitions to and from methylotrophic growth substrates, we evolved populations founded on wild-type (WT), Δ*efgA*, and Δ*ttmR* strains by alternating succinate (3.5 mM) and methanol (15 mM) as the sole carbon and energy source in minimal media (**Fig. S1**). These strains have been noted for their phenotypic differences in the capacity to switch between carbon sources. In particular, the Δ*efgA* and Δ*ttmR* strains are defective in transitioning to formaldehyde-generating growth substrates, such as the transition from succinate to methanol(20). Further, the physiological bases underlying their respective defects are distinct, and are thus expected to possess distinct selective pressures (21).

We observed that by 174 generations (29 transfers), all populations had improved significantly in their ability to oscillate between methylotrophic and non-methylotrophic growth substrates, and grew even more rapidly than the ancestral WT strain (**Fig. 1A**). A closer phenotypic examination into the transitions to methylotrophic growth revealed decreases in lag times and increases in growth rates during the succinate-to-methanol transition and decreases in both lag times and growth rates in succinate control conditions (**Fig. 1B, C**). Growth yields appeared largely unaffected by the experiment, except for lineages of the Δ*ttmR* strain, which generally suffers from a comparative growth defect that was resolved (**Fig. 1D**). In contrast, WT and the Δ*efgA* mutant lineages both experienced decreased yields in non-transition growth on succinate by generation 174 (**Fig. 1D**), suggesting a tradeoff in these features of growth in the conditions tested. Collectively, the observed evolved phenotypic changes suggest there is strong selective pressure to improve in growth measures during carbon transitions, particularly the transition to methylotrophy.

**Figure 1.**
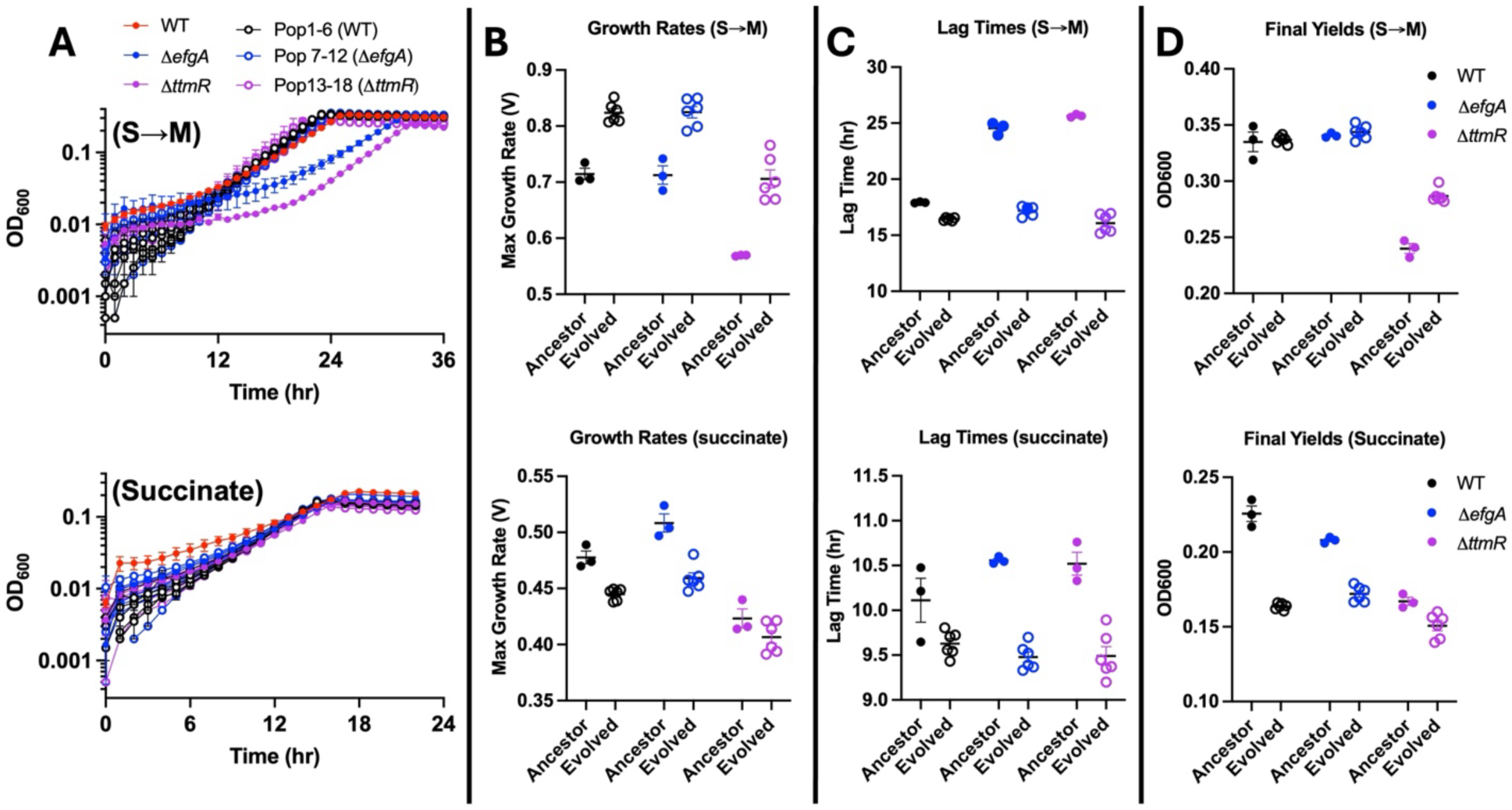
Repeated switching between methylotrophic and non-methylotrophic growth leads to rapid phenotypic improvements for evolved populations. A) Growth of evolved populations from generation 174 seeded with WT, Δ*efgA*, and Δ*ttmR* in comparison with the founding strains, either in the transition to methylotrophy (top) and in a non-transition in succinate media (bottom). Corresponding matched growth metrics of those ancestor strains and evolved populations were measured, including B) Growth rates, C) Lag times, and D) Growth yields. Ancestor strains were grown in biological triplicate, and evolved populations were grown in technical duplicate; points represent the mean and error bars represent standard error of the mean (SEM). Abbreviations: M, methanol; S, succinate

### Genetic characterization of adaptive changes identifies an uncharacterized response regulator receiver as a selective target in wild-type populations

To understand the genetic changes underpinning the phenotypic changes of WT during carbon switching, we conducted genomic resequencing of three replicate evolved populations founded on WT at generation 174 (**Table S2**). Notably, we found all six WT populations contained high frequencies of mutations to the *Mext_1965* gene locus with mutant alleles at majority frequencies, ranging from 55% to 100% (and 5 of 6 at frequencies greater than 90%) (**Table 1**). These data suggested that most populations were already swept or on a trajectory to be swept by *Mext_1965* mutants completely.

The evolved alleles of *Mext_1965* consisted of nonsynonymous base substitutions as well as nonsense and frameshift mutations, suggesting there was selection for loss-of-function mutations. In agreement with this, a deletion of the *Mext_1965* recreated the growth phenotypes that we observed for the WT populations at large, as did a representative isolate strain (JB55) that possessed the single-base deletion at position 310 of 381 annotated nucleotides (**Fig. 2A**). Curiously, all 6 WT lineages contained a subpopulation carrying this specific mutation (**Table 1**). Furthermore, the high degree of parallelism among the evolved mutations suggested a strong selection upon the *Mext_1965* locus for loss-of-function mutations in the WT background in the repetitive switching conditions tested. Indeed, when we tested a Δ*Mext_1965* mutant in competition against WT we saw that it had the strongest selective advantage under carbon switch conditions, where it was highly beneficial to transition either from succinate-to-methanol or from methanol-to-succinate (selection coefficients: succinate-to-methanol transition, S_S®M_ = 0.42, methanol-to-succinate transition, S_M®S_ = 0.56) (**Fig. 2B**). However, Δ*Mext_1965* outperformed WT under all conditions tested, even when there was no transition in the sole carbon source (selection coefficients S_S®S_ = 0.31, S_M®M_ = 0.12). Due to the overall beneficial impact of mutating *Mext_1965* on the examined carbon transition phenotypes, both in evolving populations and in direct paired competition assays, and its annotation as a response regulator, we designated this gene as encoding ***cstR***, for **<u>c</u>**arbon **<u>s</u>**ource **t**ransition **<u>r</u>**egulator.

**Figure 2.**
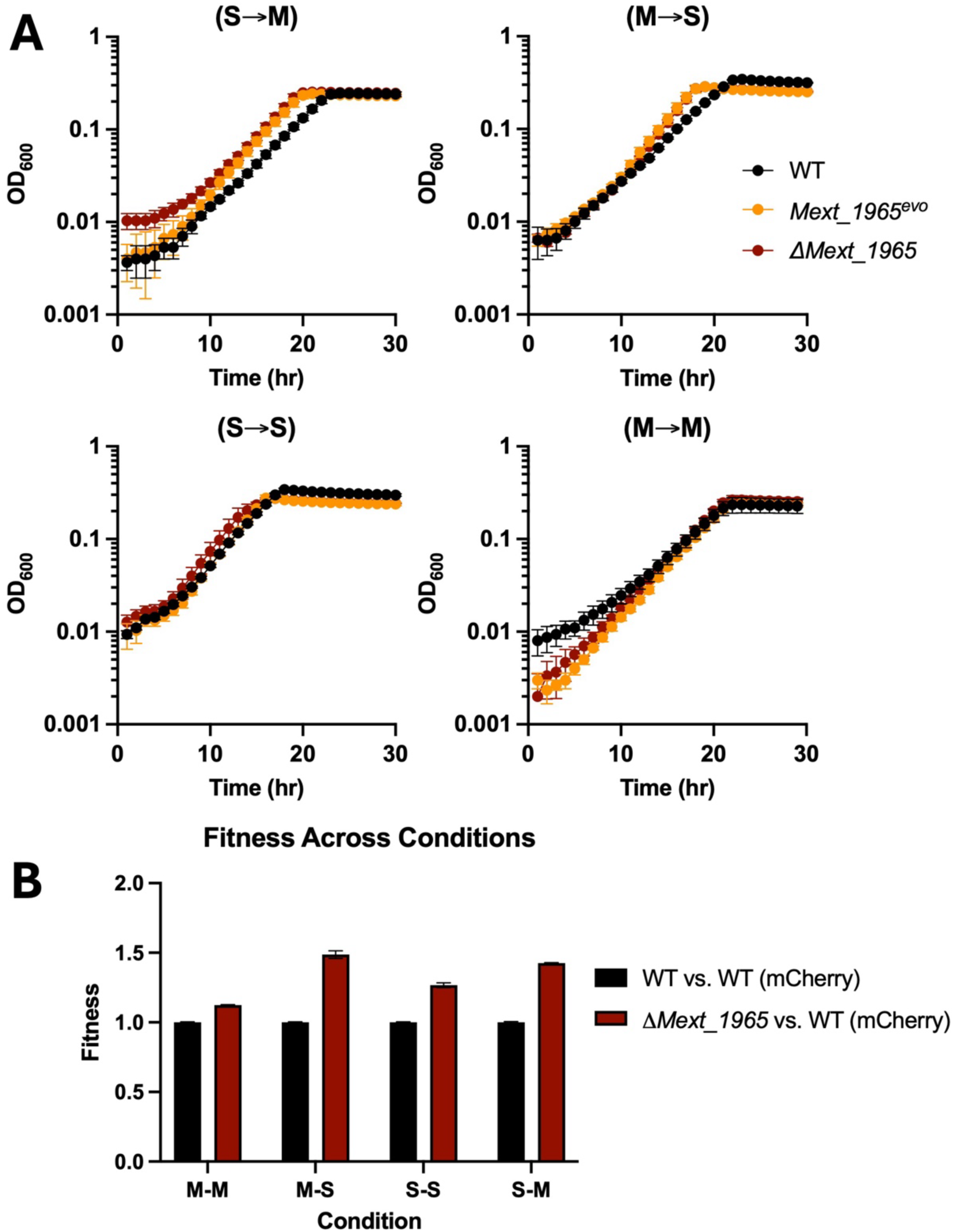
*Mext_1965*^evo^ is a loss-of-function allele that provides growth advantages over multiple conditions. A) Growth of a representative *Mext_1965^evo^* strain (JB55) isolated from a WT population containing a single fixed mutation was assessed in both transition (top) and non-transition (bottom) conditions between succinate (S) and methanol (M). A deletion strain (Δ*Mext_1965*) was also included. B) The Δ*Mext_1965* mutant was cocultured with WT in order to determine relative fitness in each of these conditions. Strains were grown in biological triplicate; points represent the mean and error bars represent standard error of the mean (SEM).

### Evolved populations of Δ*efgA* and Δ*ttmR* mutants experienced disparities between observed *cstR* mutation frequencies despite enhanced phenotypic effects

Mutations in *cstR* occurred to a much lesser extent in evolved populations at the same generational timepoint for the Δ*efgA* and Δ*ttmR* mutants. While *cstR* mutations essentially fixed in all WT populations by generation 174 (range = 55.6-100%, >83% in 5/6 populations), they were observed at intermediate frequencies (43.8-66.7%) in five of six Δ*efgA* populations and only observed in two Δ*ttmR* populations (frequencies: 7% and 56.2%) (**Table 1**). This pattern hinted at possible limitations of mutations in *cstR* in those genetic backgrounds to improve growth phenotypes during carbon source transitions. Thus, we investigated whether there were differences in the contribution of Δ*cstR* mutations in the Δ*efgA* and Δ*ttmR* strain backgrounds and found that the Δ*cstR* deletion provided a distinct growth advantage. Both resulting strains, Δ*efgA* Δ*cstR* and Δ*ttmR* Δ*cstR*, had dramatic improvements in growth during the succinate-to-methanol transition compared to the single mutant parental strains, and with no observable tradeoffs in the methanol-to-succinate transition (**Fig. 3**). Specifically, the lag defects were abolished and the growth rates improved, surpassing that of WT, and the yield defect of the Δ*ttmR* mutant was also alleviated. This contrasted with our expectations given the disparities in observed mutational frequencies to *cstR* and demonstrated that the loss of *cstR* provided striking growth advantages in all three genetic backgrounds during transitions between succinate and methanol. Taken together, these data suggest that the CstR-mediated signal transduction pathway is required to observe the deleterious effects of the Δ*efgA* and Δ*ttmR* mutations upon transitioning to methylotrophy.

**Figure 3.**
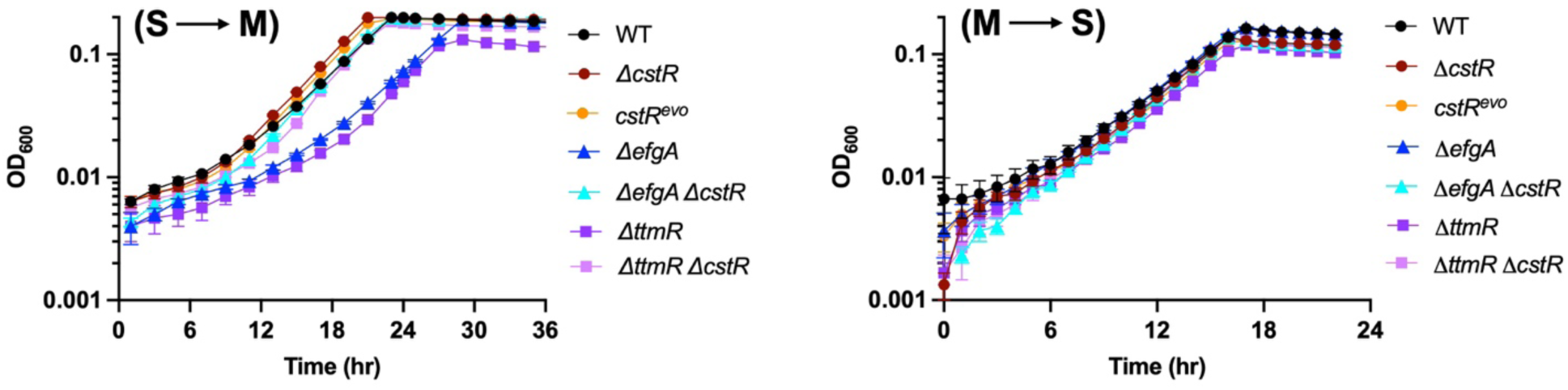
Removing *cstR* improves the transition to methylotrophy in multiple genetic backgrounds. The Δ*cstR* mutation (Δ*Mext_1965*) was introduced into the Δ*efgA* and Δ*ttmR* backgrounds and grown in transitions between succinate (S) and methanol (M) in both directions alongside WT, *cstR^evo^*, and Δ*cstR*. Strains were grown in biological triplicate; points represent the mean and error bars represent standard error of the mean (SEM).

### Loss of *cstR* decouples increased formaldehyde resistance from impaired transition to methylotrophy in the Δ*efgA* and Δ*ttmR* mutants

Previous investigations have demonstrated a tradeoff between the increased resistance of the Δ*efgA* and Δ*ttmR* mutants to exogenous formaldehyde and the ability to efficiently transition to methylotrophic metabolism, where endogenously generated formaldehyde is transiently elevated (20, 21). Therefore, we sought to examine whether the enhanced ability to transition to methylotrophic metabolism in Δ*cstR*-containing strains came at the expense of formaldehyde resistance. We examined formaldehyde-impacted growth phenotypes and established minimal inhibitory concentrations of formaldehyde (MIC_FA_) in all strains. Surprisingly, loss of *cstR* had no apparent effect upon the MIC_FA_ in any of the examined genetic backgrounds (**Fig. 4A**). Furthermore, when growth was monitored in the presence of intermediate levels (2-3 mM) of exogenous formaldehyde, we observed a modest growth advantage for the Δ*cstR* strain in comparison to WT (**Fig. 4B**). Thus, the absence of *cstR* decouples the tradeoff between efficiency of transitioning to methylotrophy and exogenous formaldehyde tolerance in strains lacking *efgA* or *ttmR*.

**Figure 4.**
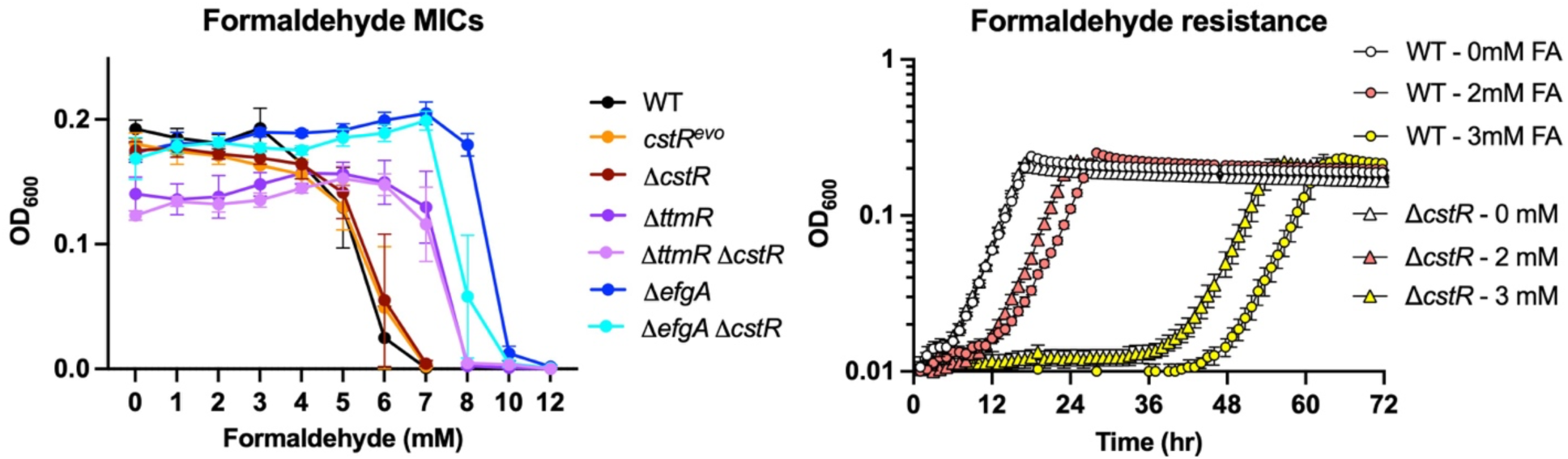
Loss of *cstR* function does not sacrifice resistance to exogenous formaldehyde. Strains were grown in MP-succinate media supplemented with a range of exogenous formaldehyde concentrations to saturation and then final OD_600_ values were measured (left). WT and Δ*cstR* were also grown and monitored in MP-succinate media supplemented with intermediate (0-3 mM) concentrations of exogenous formaldehyde (right). Strains were grown in biological triplicate; points represent the mean and error bars represent standard error of the mean (SEM).

### Examination of the novel and conserved response receiver regulator *cstR* and its global impacts

CstR is annotated as a response receiver regulator with a CheY-like characterization and is broadly conserved among the Methylobacteria clade (**Fig. S3**)(17). CheY is a canonical single-domain response regulator receiver with a key role in controlling the direction of flagellar rotation (42) but other CheY-like response regulator receivers have non-chemotactic processes (16, 43, 44). *M. extorquens* possesses four separate copies of CheY, suggesting that not all may be involved in the same aspect of chemotaxing, or involved in chemotaxis directly. Chemotaxis and cellular energetics are also intimately linked (45). When CstR was compared to protein structural databases via Gaia (33), strong matches to Rrf1 homologs existed (up to 94.4% identity), a protein first characterized in *Nitratidesulfovibrio vulgaris* Hildenborough (DVU_0530) that impacts the regulation of the high molecular weight cytochrome (*hmc*) operon whose product transfers electrons from periplasmic hydrogenases to cytoplasmic sulfate-reduction pathways (46, 47) **(Fig. S3**). Thus, in that bacterium, Rrf1 function directly impacts energy metabolism and redox balance at the membrane, suggesting a possible role for *cstR* in *M. extorquens* despite its lack of an *hmc* operon. To gain insights into the underlying physiological changes that the *cstR* mutation imparts, transcriptomic comparisons were made between the *cstR* mutant and the WT strain under succinate and succinate-to-methanol transition conditions (**Fig. 5**). Indeed, gene enrichment analysis reveals expression changes in chemotaxis/motility, signaling, and membrane genes in the Δ*cstR* mutant. In the mutant background, a significant number of differentially expressed genes (DEGs) were identified: 633 genes in the succinate condition (229 upregulated, 404 downregulated) and 306 in the succinate-to-methanol transition (89 upregulated, 217 downregulated) (**Table S3**). In succinate, there was upregulation of genes related to flagellar biosynthesis, chemotaxis-related functions, membrane composition (beta barrel proteins, LPS synthesis), nucleotide signaling (EAL- and GGDEF-domain containing enzymes), and transporters (**Table S4**). Downregulated genes in succinate include synthesis of the redox cofactor pyrroloquinoline quinone (PQQ), chemotaxis functions, transport, von Willebrand factor proteins used for adhesion, and production of Hsp20 proteins (small chaperones used to mitigate protein stress). In the transition condition, fewer functions are enriched: flagellar metabolism and chemotaxis are upregulated, while sensing (PAS-domain proteins, two component systems), cyclic di-GMP metabolism, and some chemotaxis functions are downregulated. Though genes annotated as chemotaxis-related show up across these categories, the strongest enrichment is among upregulated genes. Additionally, though not among the strongest enrichment categories, we observed downregulation of stress-related genes previously found to be strongly upregulated in formaldehyde-resistant subpopulations of the wild-type strain (48). Cumulatively, this data suggested a strong role for altered signaling and response in the Δ*cstR* strain, with a strong downstream impact on energy metabolism, movement, colonization, and stress response. When taken together, we generated a series of starting hypotheses over the function of CstR. Specifically, we predicted that CstR would impact: 1) energy metabolism and carbon utilization and transition broadly, 2) motility and colonization, or 3) tolerance and response to environmental stressors. Throughout the remainder of the manuscript, we evaluate these (not mutually exclusive) hypotheses.

**Figure 5.**
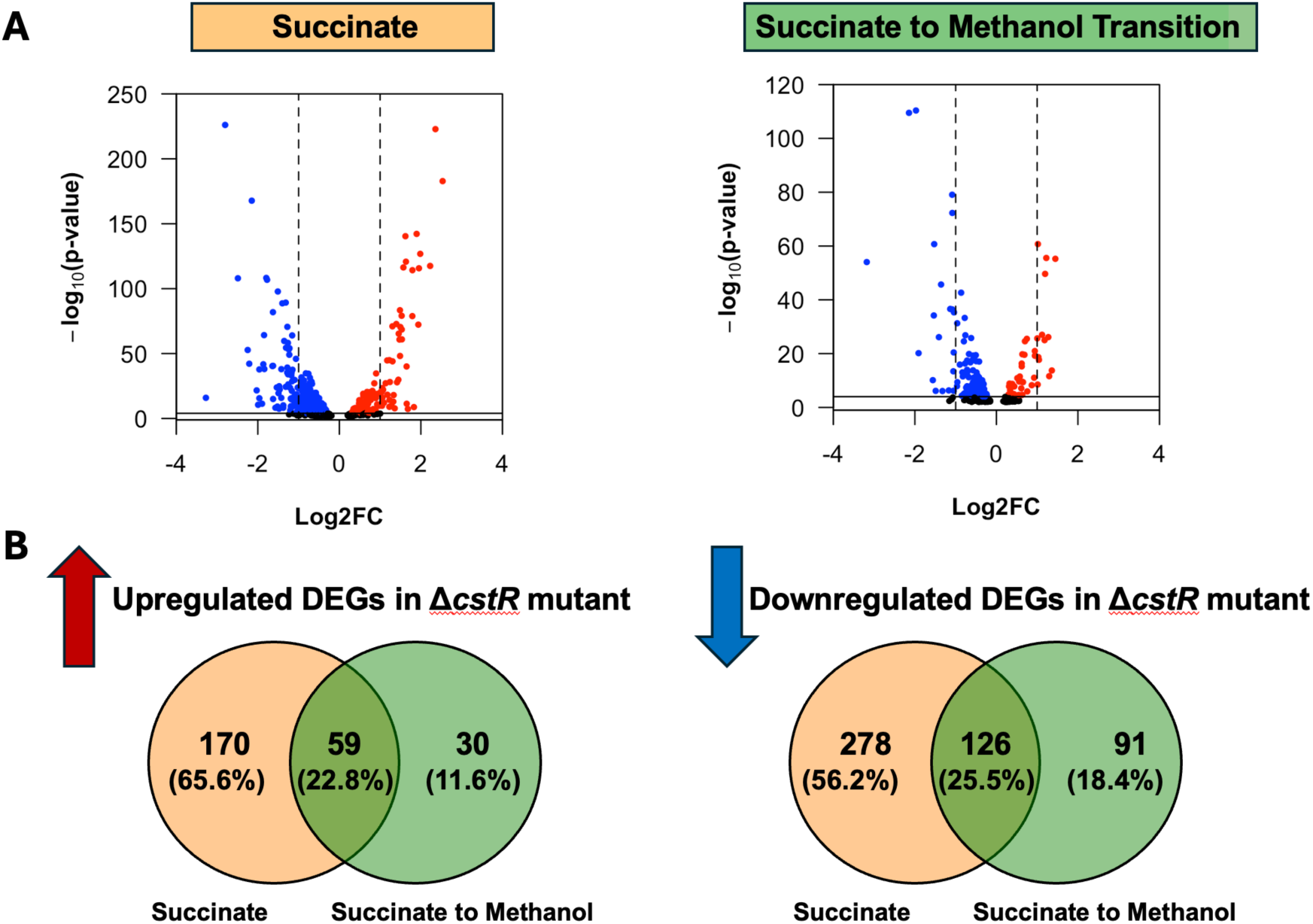
Loss of *cstR* has pleiotropic effects on the transcriptome. Gene expression was compared between WT and Δ*cstR* strains under succinate (orange) or succinate-to-methanol (green) growth conditions (biological N=3). Volcano plots (A) represent relative expression (reported as log_2_(fold-change)) against statistical confidence (reported as log_10_(p-value)). Significantly down-regulated (blue) or up-regulated (red) genes are indicated by colored dots. Dashed vertical lines indicate a |log_2_(fold-change)| threshold greater than 1. Venn diagrams (B) indicate the number and proportion of significantly up-regulated (left) and down-regulated (right) genes that are differentially expressed in succinate growth, the succinate-to-methanol transition, or both (intersection).

### Motility assays show that Δ*cstR* has increased population dispersal in succinate conditions

Due to the CheY-like characterization of CstR and the differential expression of multiple genes with predicted annotations related to chemotaxis and/or flagellar formation and motility, we sought to directly assess the impact of *cstR* upon population-level motility. We grew Δ*cstR* in several carbon sources and then evaluated their population dispersal area on succinate-containing media to ensure uniform biomass. We found that the mutant strain had increased dispersal when cells had been grown on succinate media prior to inoculation into motility media (**Fig. 6**). These data are inconsistent with CstR functioning as a canonical CheY, as a *cheY* mutant would typically be unable to chemotax through agar due to defective directional switching. Therefore, the increased dispersal area of the Δ*cstR* mutant may be unrelated to motility and instead reflect increased growth rate and/or reduced adhesion, both of which are consistent with our growth (growth rate) and transcriptomic (decreased expression of adhesion proteins) data.

**Figure 6.**
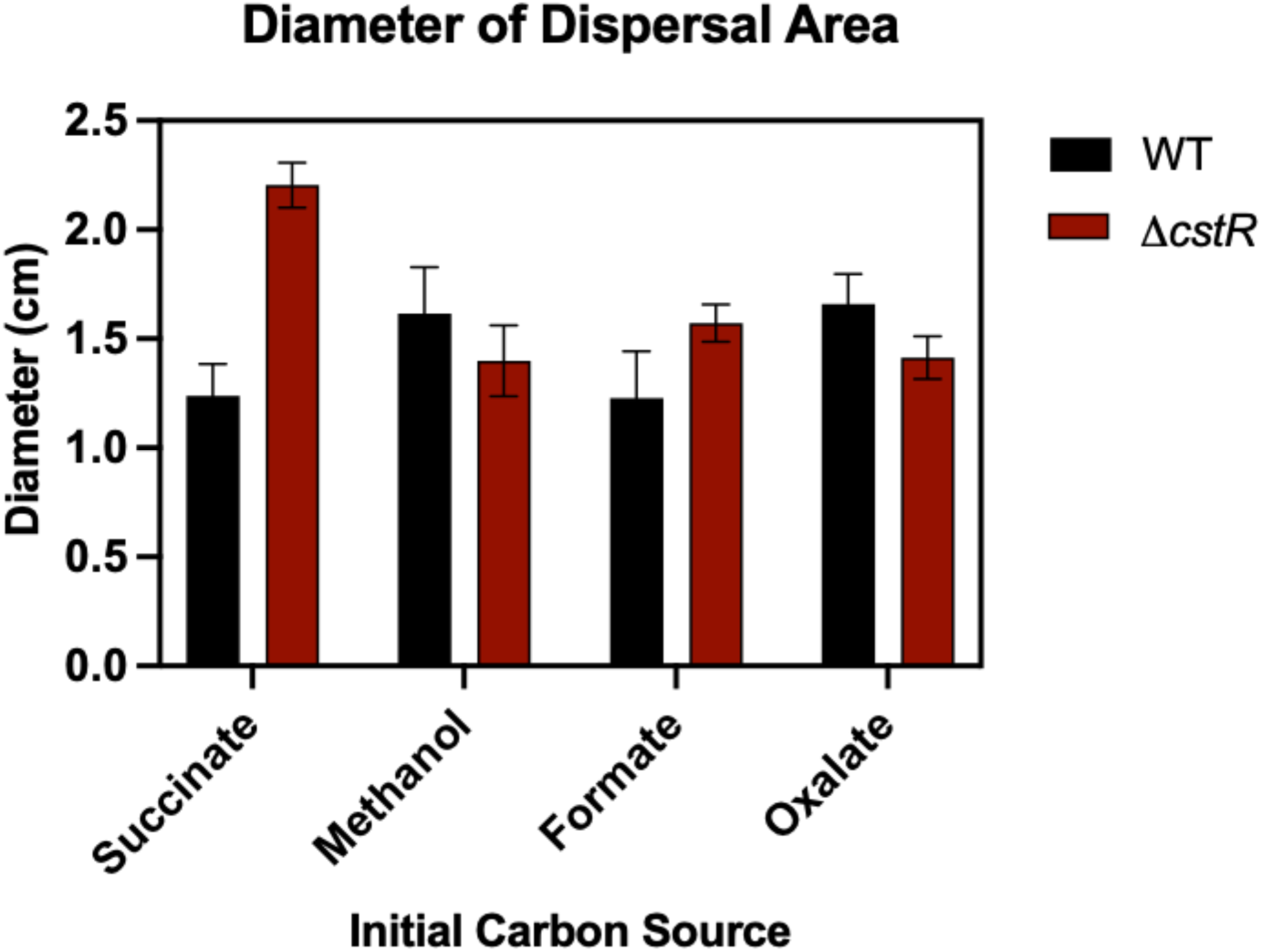
Population motility is impacted by *cstR* in succinate media. Measurement of colony diameter (cm) in motility media (0.25% agar) after completion of growth for cultures pre-grown in several separate sole carbon sources. Total colony area for three biological replicates is displayed with error bars representing standard error of the mean (SEM).

### Loss of *cstR* sensitizes cells to a subset of stressors: oxidative stress, heat, and desiccation

Given the upregulation of stress-related genes and the fact that increased motility can also be a response to stress induction, we examined the impact of deleting *cstR* under a panel of stress conditions.

Specifically, we chose elevated concentrations of ethanol, sucrose, sodium chloride, and hydrogen peroxide as well as exposure to UV radiation, heat stress, and desiccation. We found that there were no significant phenotypic effects under exposure to several stressors, and thus no universal detrimental impact of deleting *cstR* (**Fig. S4**). However, differential growth impacts were observed under heat, oxidative, and desiccation stress. Specifically, the Δ*cstR* strain demonstrated an increased loss of viability compared to the WT upon exposure to 50 ℃ (**Fig. 7A**) as well as after extended desiccation stress (**Fig. 7B**). Lastly, exposure to the oxidant hydrogen peroxide led to a demonstrable growth defect compared to WT (**Fig. 7C**). Interestingly, the transition to methylotrophy appeared to allow better tolerance to peroxide stress than continued growth on a particular single carbon source (**Fig. 7C**). The combined results from our stress panel demonstrated that there were few conditions in which loss of *cstR* function was detrimental and no conditions in which it was beneficial. Though uncertain what cellular components are more susceptible to stress in the Δ*cstR* strain, it is notable that all reported stressors in **Figure 7** have the potential to impact the cellular membrane.

**Figure 7.**
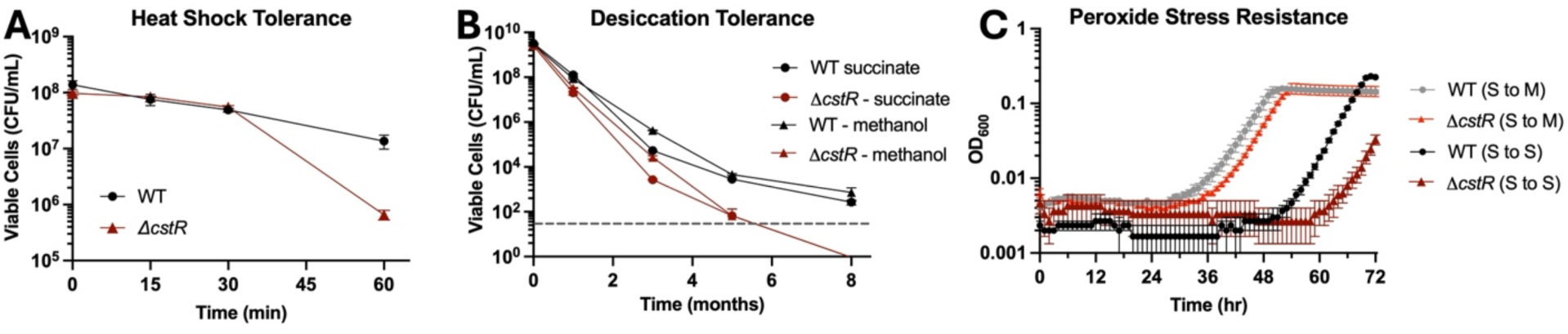
Cells lacking CstR experience heightened sensitivity to multiple stressors. WT and Δ*cstR* strains were exposed to A) extended temperature stress (survival), B) different time exposure to desiccation stress (survival), and C) 0.1% hydrogen peroxide concentrations (growth). Strains were treated in biological triplicate; points represent the mean and error bars represent standard error of the mean (SEM).

### CstR impacts cellular energy and redox balance

Cellular energetics are reliant on the coupling of redox reactions and proton gradients, whereby electron transfer from cofactors such as NADH to components of the electron transport chain (ETC) drives proton pumping, establishing the electrochemical gradient that produces ATP. We probed the *cstR* mutant using a three-pronged approach, measuring the NAD+/NADH ratio, total ATP levels, and the cellular response to extreme pH.

Cells were grown and harvested under different carbon source regimes to assess the NAD+/NADH ratio and relative ATP levels between strains and across growth phases. Specifically, we measured metabolites in early log phase and early stationary phase during growth on succinate, during the transition from succinate to methanol, and also during the transition from succinate to oxalate. We included the oxalate condition as we had previously noted that the Δ*cstR* mutant strain experiences a more rapid die-off in stationary phase in oxalate compared to WT in this condition; thus, it represented a vulnerability in the mutant strain that might prove insightful (**Fig. S5**). We observed that the Δ*cstR* mutant NAD+/NADH ratio was decreased compared to that of WT in all but one of six conditions, with only oxalate stationary phase samples having near statistical significance (adjusted P=0.0565) (**Fig. 8A**). With regard to ATP levels, no differences were observed between WT and the Δ*cstR* mutant strain in most conditions (**Fig. 8B**). The single divergence existed between the strains that were grown in oxalate as they were entering stationary phase (prior to any loss of viability). Here we found that the mutant had significantly decreased ATP levels, exposing an energetic deficit in this specific condition. Further investigation found that as oxalate is consumed in this condition, the culture media pH values dramatically rise, unlike succinate- or methanol-based media (**Fig. S5**). This suggested a connection between pH levels encountered and the ability to generate and maintain energy.

**Figure 8.**
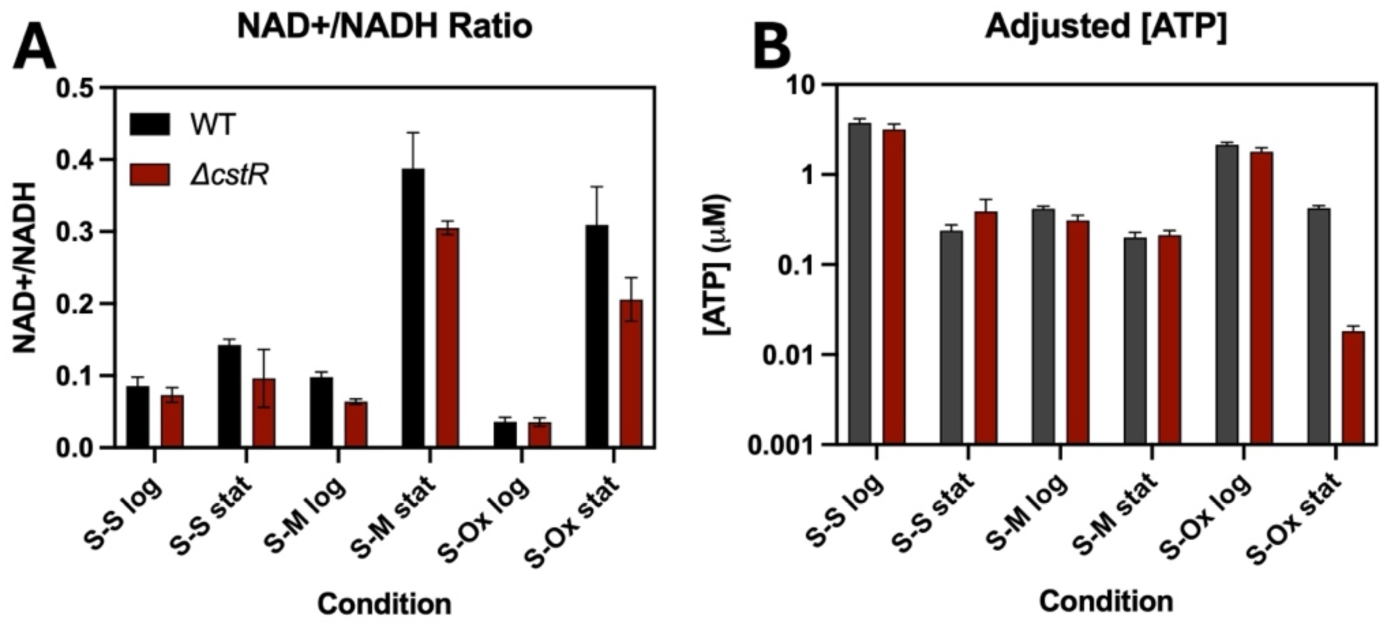
Cells lacking *cstR* have perturbed metabolite pools. The WT and Δ*cstR* strains were grown in MP media with different sole carbon sources and harvested during exponential or stationary phases of growth. Cell pellets were extracted and treated to assay conditions allowing luciferase-linked detection of ratio of oxidized and reduced nicotinamide adenine dinucleotide (NAD+/NADH, left) and concentrations of adenosine triphosphate (ATP, right). Bars represent mean values, error bars depict SEM, biological replication, N=3-4. S: succinate, M: methanol, Ox: oxalate, log: logarithmic/exponential phase, stat: stationary phase.

We further probed the Δ*cstR* mutant’s growth and survival under pH stress, both acidic and alkaline, during growth on succinate and survival once carbon is exhausted. In alkaline conditions, the mutant maintained growth and survival comparable to WT until pH 10 where the Δ*cstR* mutant had a growth defect and decreased survival; this is comparable to pH levels observed in oxalate media linked to increased mortality of the mutant strain (**Fig. 9A, B**). In acidic conditions, the Δ*cstR* mutant grew better than WT and had identical survival rates (**Fig. 9C, D**). These data suggest that the Δ*cstR* mutant has an altered internal pH and corresponding sensitivity to alkaline external pH.

**Figure 9.**
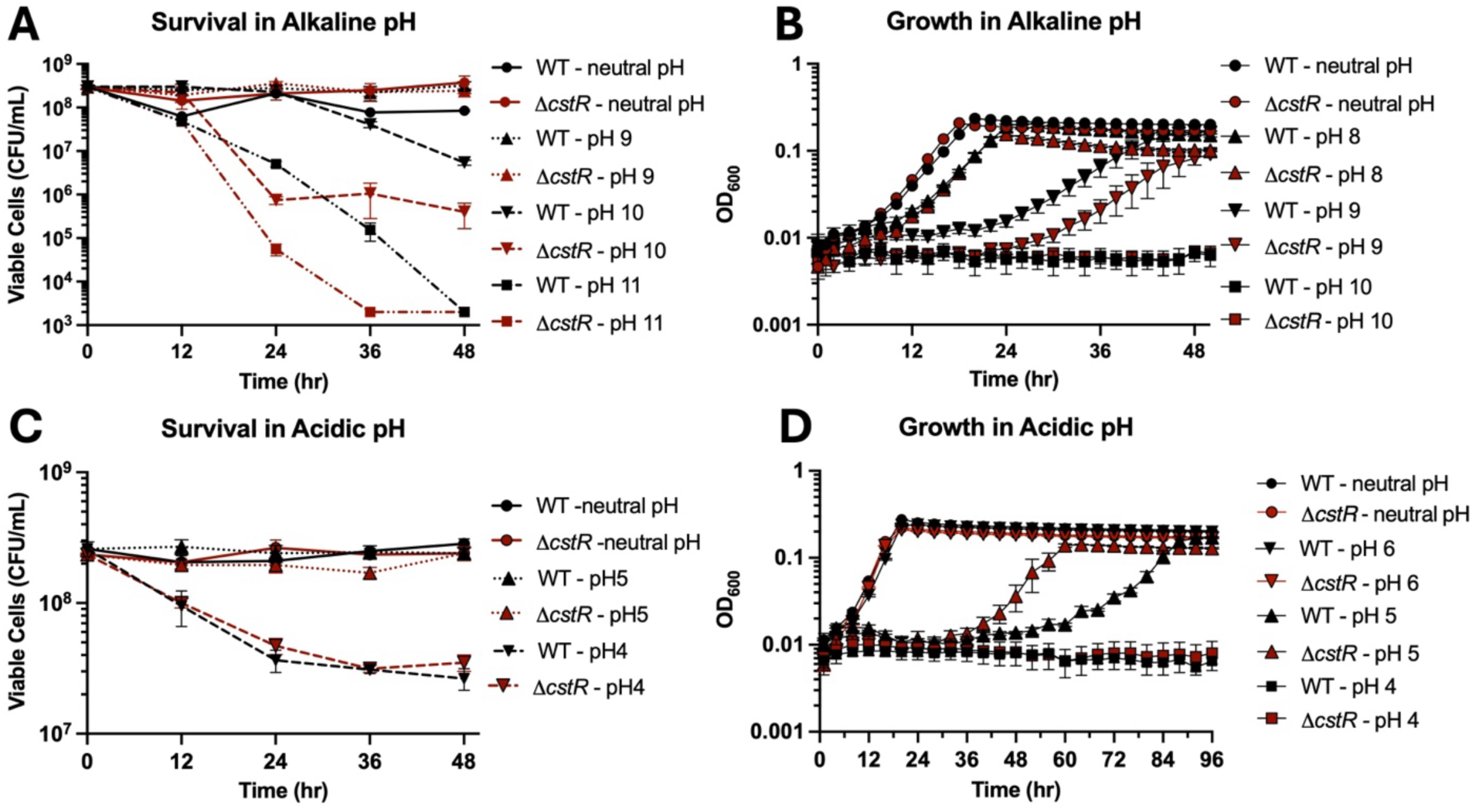
CstR function impacts growth and survival under altered pH conditions. WT and Δ*cstR* strains were grown to stationary phase in MP-succinate media and exposed to a range of alkaline (A, B) or acidic (C, D) pH conditions. Survival over time under different pH conditions (A, C) were conducted by resuspending cells in carbon-less MP media and conducting time lapse dilution plating to obtain viable cell counts. Growth under different pH conditions (B, D) was conducted by diluting into MP-succinate media and monitoring OD_600_ over time. Plotted values represent a mean of three biological replicates, and error is represented by SEM.

Collectively, these data suggested that the Δ*cstR* mutant has an elevated NADH pool compared to WT. NADH can serve as a supply of electrons with which to generate proton motive force (pmf) and ATP generation during conditions conducive to cell growth and division; however, accumulation of NADH may also indicate a bottleneck in the respiratory chain/process. Thus, at low or neutral pH, when the extracellular environment is saturated with protons and the cell is attempting to maintain homeostasis by pumping even more protons out of the cell, elevated NADH could beneficially support respiration and helps to drive this momentum. Conversely, at high pH, when the extracellular environment has reduced protons, outcomes will depend more on the cell’s capacity to generate sufficient pmf against a proton-poor exterior. The lethal effects observed in both strains under alkaline conditions, but exacerbated in ΔcstR, result from systemic failures in pH homeostasis or redox balance. Altogether, this suggests that the loss of CstR leads to imbalances in the ability of the cells to balance their redox pools and generate energy.

### Loss of CstR does not uniformly improve defective transitions to methanol-based growth

We previously found that, compared to the transition from succinate to methanol, cells lacking EfgA and TtmR had an exacerbated transition defect from formate- to methanol-based growth. We sought to determine whether loss of *cstR* would also be beneficial under this transition, as formate is another available carbon source in the phyllosphere, and this transition has been previously observed to impart a more dramatic growth defect (21).

When exposed to this alternative transition, we found that the *cstR* deletion differently impacted each genetic background: it was i) again slightly beneficial in the WT background (though not with statistical significance), ii) neutral in the Δ*efgA* background, and iii) deleterious in the Δ*ttmR* background, leading to an exacerbated growth defect in the form of a further extended lag phase (**Fig. 10**). Therefore, the formate-to-methanol transition reveals distinct epistatic interactions between these alleles and the *cstR* deletion. These data also suggest that the regulation provided by TtmR and CstR is complementary and potentially redundant, and that loss of their collective regulation impacts an exacerbated defect during the formate-to-methanol transition, a C_1_-to-C_1_ transition.

**Figure 10.**
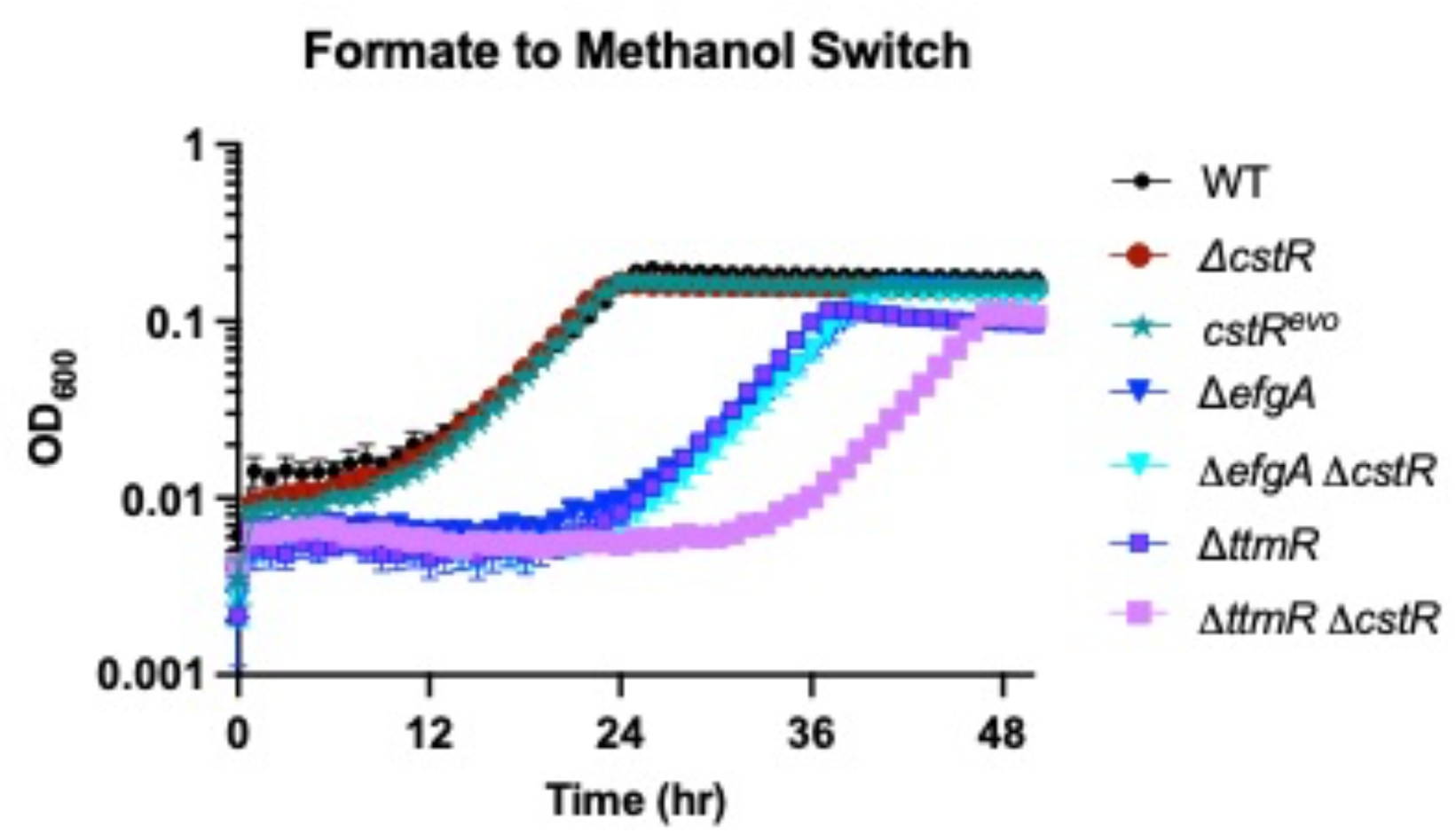
CstR provides genotype-specific effects in a formate-to-methanol transition to methylotrophy. The Δ*cstR* mutation was introduced into the Δ*efgA* and Δ*ttmR* backgrounds and grown in a transition from formate to methanol and OD_600_ was monitored to track growth over time. Strains were grown in biological sextuplicate; points represent the mean and error bars represent SEM.

The defect of the Δ*ttmR* mutant during the formate-to-methanol transition provided an optimal growth condition to test the complementation of *cstR* gene function, gauged by restoration of the decreased lag phase in the transition. When *cstR* was introduced and expressed ectopically via plasmid into the Δ*ttmR* Δ*cstR* double mutant strain, complementation was provided (**Fig. S6**). Observed complementation occurred for both the primary annotation of the *cstR* CDS (provided on pEB200) as well as an extended ORF of the CDS with an alternative start codon that cumulatively contained an additional 162 upstream nucleotides (corresponding to 54 additional codons) from the predicted translational start site (provided on pEB201). Conversely, plasmid expressing a reverse transcribed ORF contained within the annotated genomic CDS that would also be impacted by the *cstR^evo^*mutation (provided on pEB202) did not provide complementation, demonstrating that this product was not the causative factor leading to our observed phenotypic changes upon deletion from the genome. The experiment demonstrated that the 381-nucleotide annotated ORF is sufficient to provide proper complementation of function.

## DISCUSSION

In this work, we have identified a previously uncharacterized single-domain response regulator (SD-RR), CstR, in *Methylobacterium extorquens* PA1 that confers numerous, largely subtle, phenotypes to cells when its function is removed. Positive physiological changes result under more permissive conditions, including more rapid growth under exposure to novel carbon source conditions due to higher levels of reducing power. This leaves mutant cells more susceptible to damage and mortality under exposure to an array of stressors that include heat shock, desiccation stress, reactive oxygen species including peroxide, and alkaline shifts in environmental pH levels (**Fig. 11**). Overall, *cstR* loss-of-function mutants appear to be primed for more rapid growth, but simultaneously more susceptible against stress exposure; thus, this CstR protein is nested in the cellular signaling/regulatory network that allows cells to successfully navigate environmental shifts and make decisions on when to invest in growth/reproduction versus maintenance/protection. The lack of stark phenotypes may be a result of CstR optimizing metabolism, possibly through multiple inputs, within a highly branched and interconnected regulatory network (**Fig. 11**).

**Figure 11.**
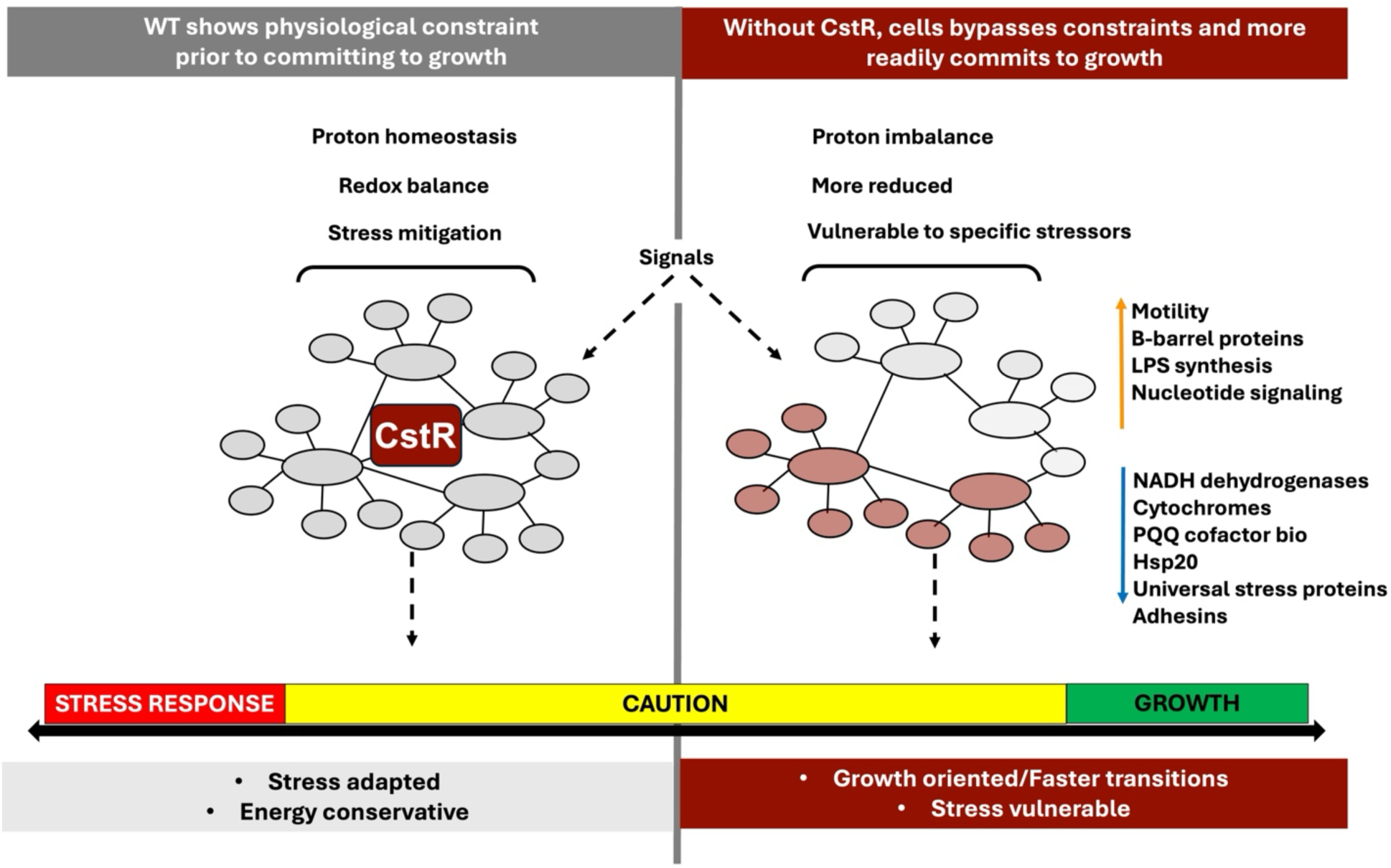
Model depicting cellular impacts imposed by the presence (left) or absence (right) of CstR in *Methylobacterium extorquens* PA1.

Transitioning between distinct carbon substrates requires substantial remodeling of the cellular metabolic network and is likely a frequent occurrence for methylotrophic bacteria inhabiting the phyllosphere. We previously demonstrated that the model methylotroph and competitive phyllosphere colonizer *M. extorquens* PA1 exhibits a tradeoff between heightened resistance to exogenous formaldehyde and growth delays upon the transition to methylotrophy, driven by heightened levels of endogenously generated formaldehyde (20, 21, 29, 49). Therefore, it was especially interesting that populations from all three included genetic backgrounds quickly achieved dramatic improvement and indistinguishable growth in the transition from succinate to methanol, despite very different starting capacities to do so. We initially hypothesized that all populations might have adapted through identical mechanistic means; however, after pursuing sequencing of evolved populations at the *cstR* locus, we determined that this was not the case.

We intriguingly found that, in the WT background, repeated transitions between the organic acid succinate and the volatile C_1_ compound methanol selected for parallel mutational changes to the *Mext_1965* locus, now renamed *cstR*, which encodes an SD-RR. Previously uncharacterized in *M. extorquens*, follow-up work revealed that the deletion of *cstR* recapitulated growth transition phenotypes of the evolved alleles, suggesting that the predominantly selected gene variants encoded for a loss of gene function. Notably, no detected mutations affected any of the 10 aspartate sites of CstR (7.9% by composition), to potentially alter signaling transmission of the protein, which could indicate that disrupting a single phosphorylation site would be insufficient to achieve this phenotype. Additional experiments upon different carbon sources demonstrated a broad improvement in growth across tested environments, including in the presence of supplemented formaldehyde (**Fig. 4**). It was interesting however, that the loss of the *cstR* also improved growth of the Δ*efgA* and Δ*ttmR* strains, despite evolved alleles occurring with far less frequency in these genetic backgrounds (**Table 1**).

In other Alphaproteobacteria, SD-RRs have been found to act as hubs between stress response and growth, a primary example being MrrA in *Caulobacter crescentus* (50). MrrA accepts phosphate groups from numerous histidine kinases and mediates the decision between the general stress response and cell division. In this context, the deletion of *mrrA* led to an inability to activate the general stress response and failure to manage certain stressors, including those encountered during stationary phase. Many response regulators, and many SD-RRs are encoded in the genomes of environmental Alphaproteobacteria such as *M. extorquens* (17), likely out of the need to sense and respond to myriad environmental cues to conditions populations of cells encounter. This provides an expansive set of mutational opportunities to adaptively alter the cell’s existing regulatory circuitry. Consistent with this, the differences in selection for loss of *cstR* function and environment-dependent impact suggested very different regulatory configurations among the strains tested. An interesting impact of *cstR* LOF was strong downregulation of the dicarboxylate transporter *dctA* (*Mext_3076*), with impacts on the uptake of C_4_ compounds like succinate and fumarate. This could indicate repression due to carryover compounds when samples were diluted in transition experiments.

The Δ*cstR* mutation provided a strong beneficial impact under the selective pressure of repetitive carbon switching, but we postulated that there would exist condition(s) where loss-of-function would be costly, potentially under persistent stress or during transitions between other carbon sources, as such conditions can also be accompanied with inherent stress, such as when cells must newly navigate endogenously generated toxic molecules like formaldehyde. Notable to this point, the genetic loss of *cstR* function does not universally render cells vulnerable to stressors, but rather, specific stress phenotypes are affected (heat, desiccation, peroxide stress) and are thus likely conferred by the particular molecular and mechanistic changes underlying the altered growth phenotype, including altered redox and proton homeostasis. Further, the loss of *cstR* does not universally aid in the improvement of transitioning between all carbon substrates. When a transition from formate to methanol was tested, deleting *cstR* provided very different fitness impacts to the different genetic backgrounds tested: still providing a slight beneficial effect to WT, a neutral outcome to the Δ*efgA* mutant, and a detrimental outcome to the Δ*ttmR* mutant (**Fig. 10**). This demonstrates that the formate-to-methanol transition i) is more stressful to *M. extorquens*, likely a function of the relatively lower energy and redox contributions provided by formate, ii) provides a different selective environment compared to the succinate-to-methanol transition, and iii) reveals epistasis between *cstR* and the particular strain’s genetic background. Interestingly, the relative strain-level impacts of deleting *cstR* to the formate-to-methanol transition also follow the same order of frequencies of *cstR* loss-of-function mutations detected in the evolved populations (**Table 1**).

Though we expected a stronger selection to be provided by the succinate-to-methanol transition due to initial delays of the Δ*efgA* and Δ*ttmR* strains because of perturbed formaldehyde homeostasis(20, 21), some of the phenotypes of the Δ*cstR* mutant were most pronounced in the succinate growth condition. These phenotypes included the sensitivity to peroxide stress (**Fig. 7**) and the colony dispersal area when inoculated into motility plates (**Fig. 6**). Interestingly, both strains handle peroxide stress better in the succinate-to-methanol switch condition (**Fig. 7**). This indicated that some aspect of the transition to methylotrophy contributed a cross-benefit to the tolerance and growth under oxidative stress, possibly priming cells for mitigation of this stress.

Growth on oxalate was originally pursued to evaluate an electron-poor carbon source known to be available on plant leaves (51–53). We observed a large increase in pH over the course of growth of cultures in oxalate media (**Fig. S5**), a pattern consistent with previous observations of alkalinization resulting from bacterial oxalate metabolism (54–58). Oxalate is transported into *M. extorquens* cells via an oxalate/formate antiporter (59) and is oxidized to form CO_2_, thereby contributing to the rise in environmental pH via associated carbonate formation. The observation that transhydrogenase is highly upregulated in closely related *M. extorquens* AM1 cells grown on oxalate further suggests a dysregulated redox state that requires modification to achieve homeostasis (59). When grown in oxalate-containing media, the Δ*cstR* mutant strain experienced a dramatic loss of viability upon entering stationary phase, associated with heightened pH levels (**Fig. S5**). Indeed, independent of oxalate, we largely observed increased mortality and growth inhibition for Δ*cstR* populations exposed to alkaline pH conditions (**Fig. 9**). Together, this suggests a pattern of accelerated short-term growth at the cost of long-term survival in conditions where cells might experience fluctuating environmental conditions, including shifts in carbon source availability. Thus, the CstR regulator acts to provide a cellular balance between growth on different carbon sources and transitions between them, and in particular the balance between growth and survival.

Collectively, it appears that removing CstR-mediated signaling and regulation alters cellular redox and intracellular pH that leads to a steeper proton gradient across the membrane, thereby increasing the proton motive force under neutral media conditions and advantageous transitions between succinate and methanol. This may also underpin the observed motility phenotype of the mutant strain in succinate, as flagellar rotations is powered by the pmf. However, this comes at a cost under certain stressors, including oxidative stress and alkaline pH conditions. The altered physiological state provoked by eliminating CstR allows cells to navigate transitions into new resource environments and onto novel carbon sources under neutral pH conditions, but also with potential consequences in stressors experienced in the phyllosphere as well as to cells not in a state of active growth.

## Acknowledgements

We thank Kathryn R. Fixen for reviewing the manuscript and providing insightful feedback. This work is funded by NIH NIGMS 1R35GM146904. The authors declare no conflicts of interest.

## Data accessibility statement

RNAseq data is available on the NCBI Gene Expression Omnibus (GEO). Genome sequencing files and amplicon data are available online on the NCBI Short Read Archive (SRA).

---

**Figure S1.**
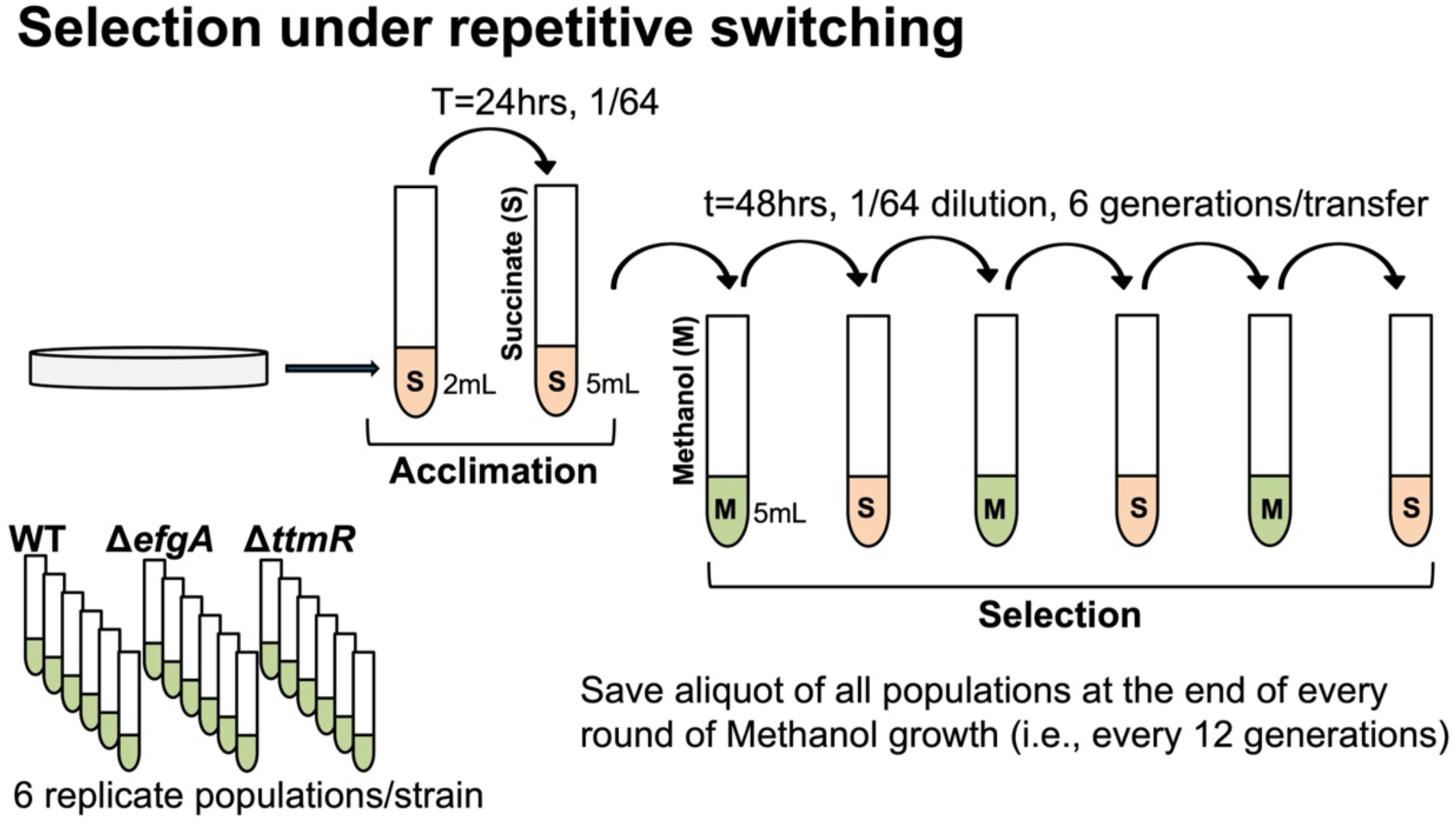
Selection regime of *M. extorquens* switching to and from methylotrophic growth. *M. extorquens* was experimentally evolved in MP liquid media for 174 generations (29 transfers). For each of three strains used (WT, Δ*efgA*, Δ*ttmR*), six independent replicate populations were used. Initial growth conditions relied on 3.5 mM succinate as a sole source of carbon and energy to acclimate cells to this mode of growth. At the first transfer, stationary phase cells were subcultured (1/64) into fresh media containing 15 mM methanol as a sole source of carbon and energy, transitioning their metabolic mode to methylotrophy. Populations were then continuously transferred between succinate and methanol-based growth at 48-72 hr intervals (approximately 6 generations). The resulting populations were sampled and saved periodically to assess genetic and phenotypic traits. Abbreviations: M, methanol; S, succinate.

**Figure S2.**
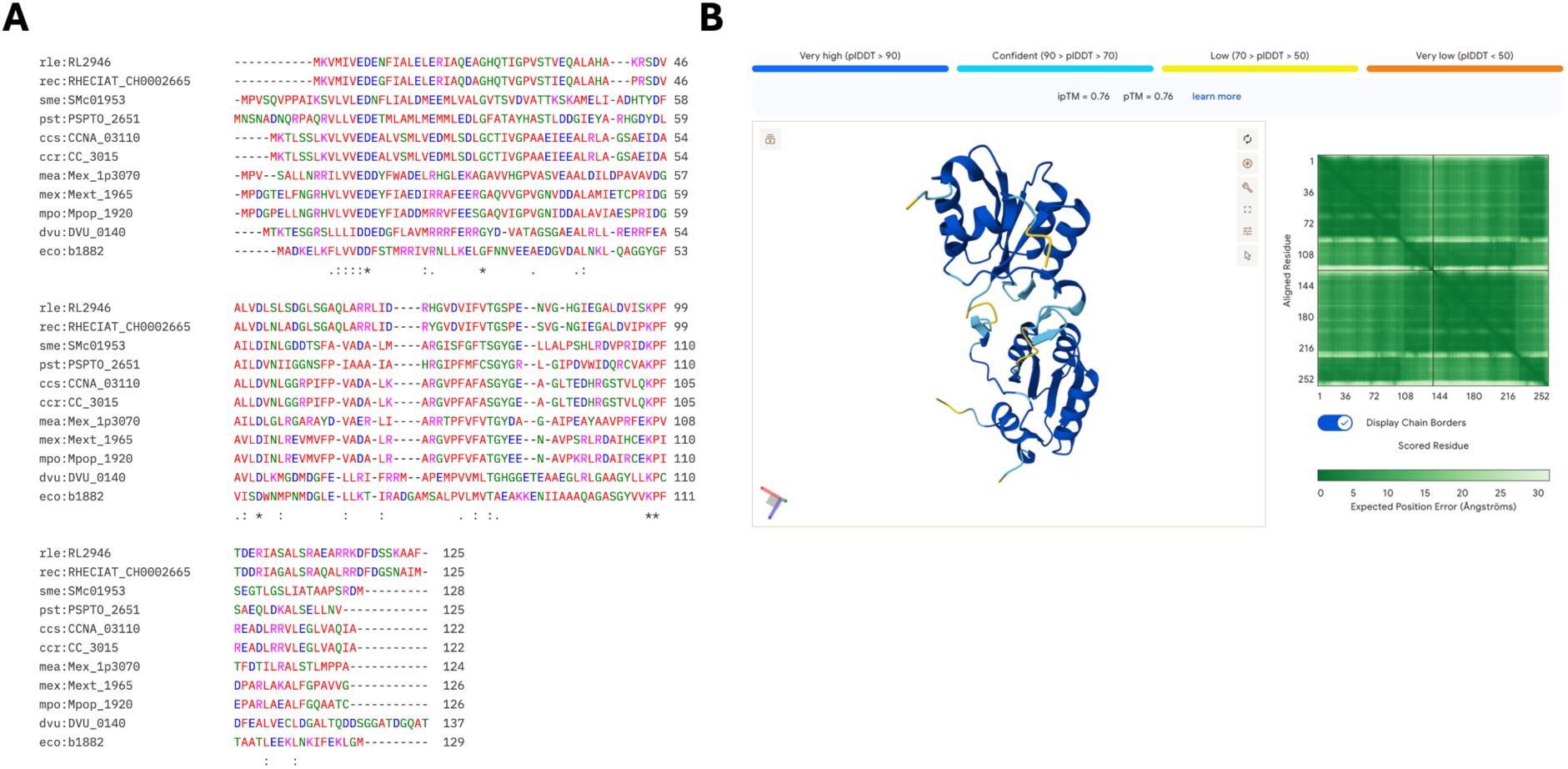
Alignments of the CstR protein with homologs. A) A Clustal Omega (26) generated multiple sequence alignment of CstR from *M. extorquens* PA1 (mex) and high similarity homologs from *M. extorquens* AM1 (mea), *M. populi* (mpo), *Caulobacter vibrioides* NA1000 (ccs), *Caulobacter vibrioides* CB15 (ccr) MrrA protein, *E. coli* K-12 MG1655 (eco) CheY protein, *Rhizobium johnstonii* 3841 (rle), *Rhizobium etli* CIAT 652 (rec), *Sinorhizobium meliloti* 1021 (sme), *Pseudomonas syringae* pv. tomato DC3000 (pst), *Nitratidesulfovibrio* (formerly *Desulfovibrio*) *vulgaris* Hildenborough (dvu) Rrf1 protein, and a high similarity Rrf1 homolog from *Methylobacterium* sp001542815. Conservation of residues is indicated when identical (*), strongly similar (:), or weakly similar (.). Small-hydrophobic residues (less Y) are in red (AVFPMILW), acidic residues are in blue (DE), basic residues are in magenta (RHK), and hydroxyl + sulfhydryl + amine + G residues are in green (STYHCNGQ). B) A structural model of CstR predicted with AlphaFold3. Displayed is a picture of the best scored model generated.

**Figure S3.**
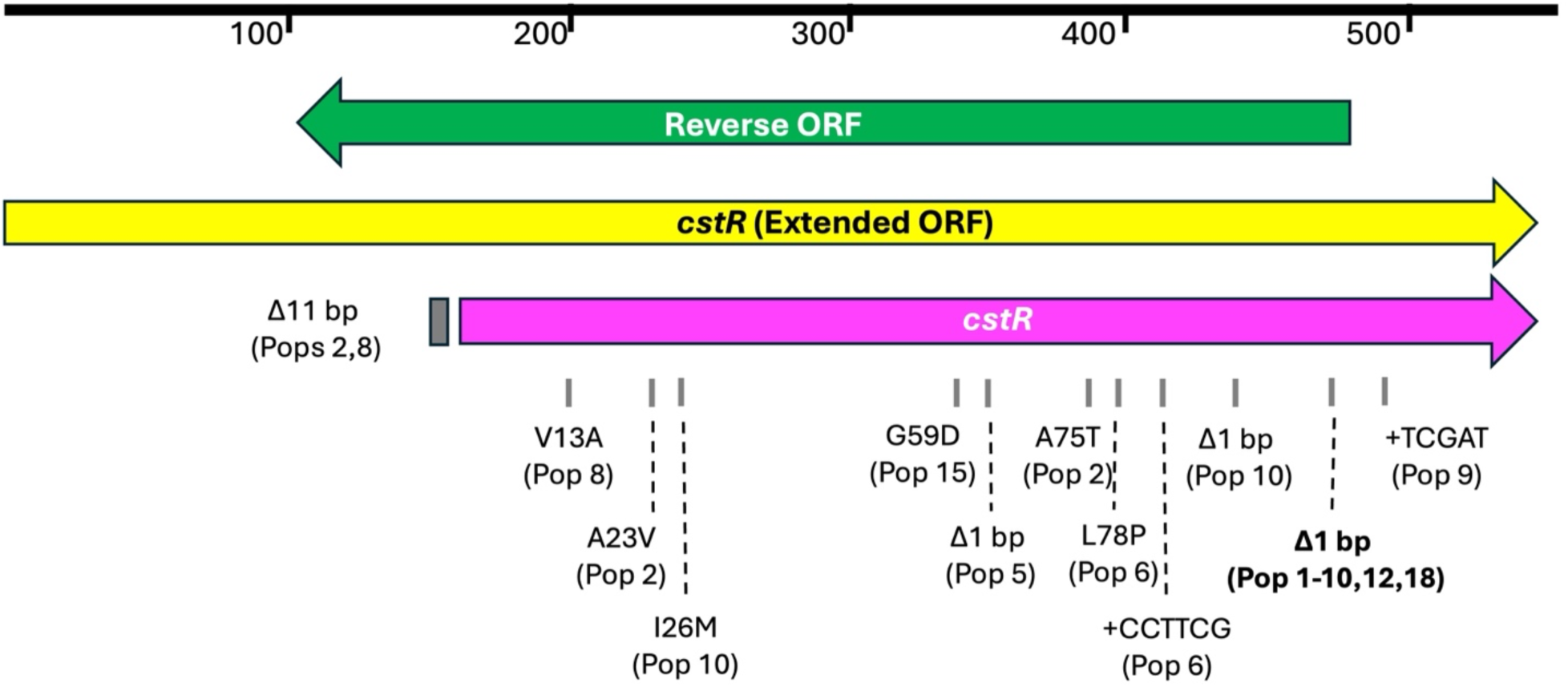
Genomic region near *cstR*. The genomic region containing *cstR* and nearby sequence is shown, with the pink arrow depicting the annotated gene, the yellow arrow depicting the extended *cstR* ORF that contains an additional 162 bp upstream of the annotated *cstR* start codon, and the green arrow depicting an ORF oriented antiparallel but overlapping a large portion of *cstR*. Gray boxes and dashes indicate the location of identified *cstR* mutations among the experimentally evolved populations (see **Table 1**). The mutation referred to throughout the text as *cstR^evo^* is indicated in bold.

**Figure S4.**
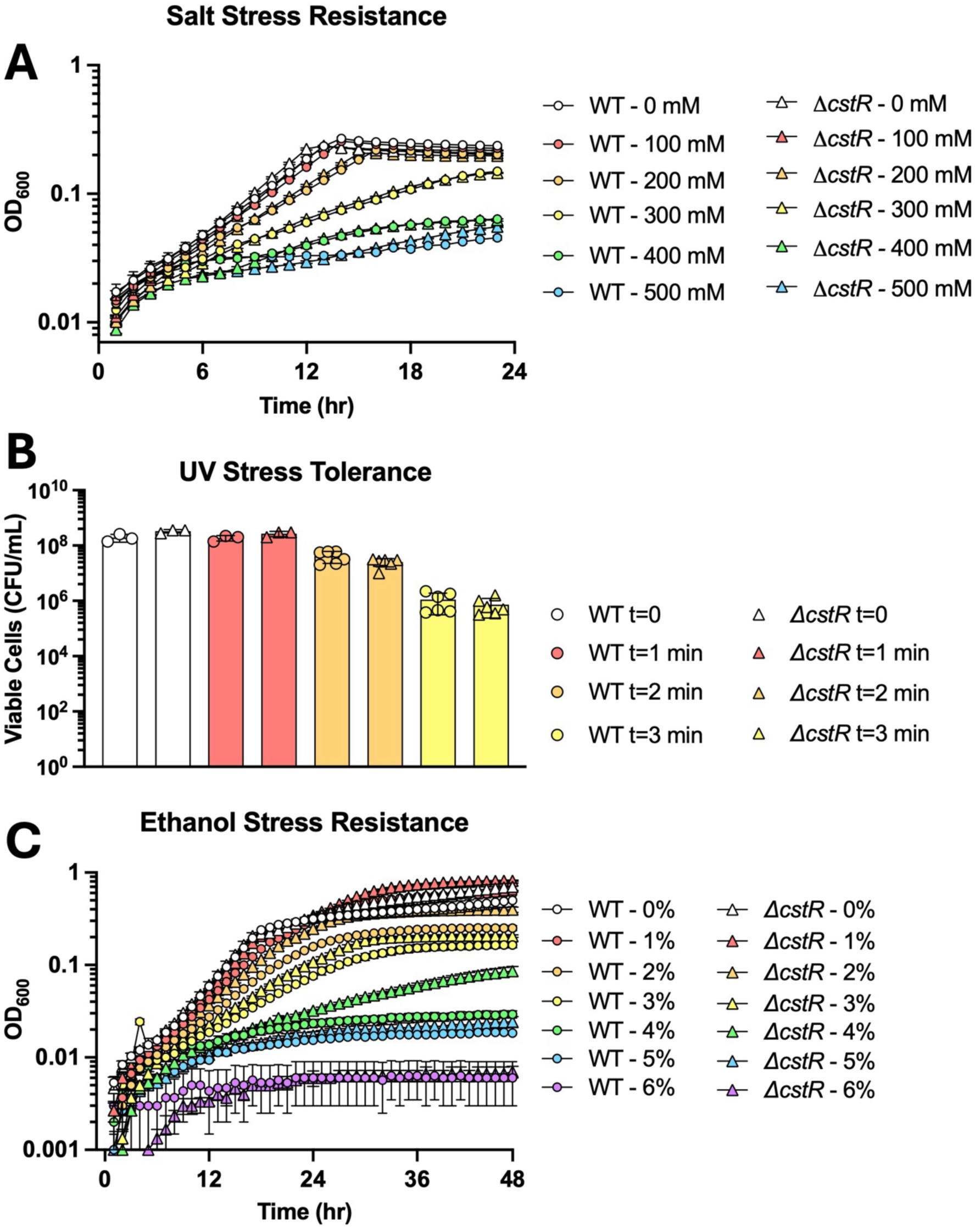
CstR does not impact resistance/tolerance to all stressors. WT and Δ*cstR* strains were exposed to A) a range of salt (sodium chloride) concentrations (growth), B) different time exposure to ultraviolet radiation (survival), and C) a range of ethanol concentrations (growth). Strains were treated in biological triplicate; points represent the mean and error bars represent standard error of the mean (SEM).

**Figure S5.**
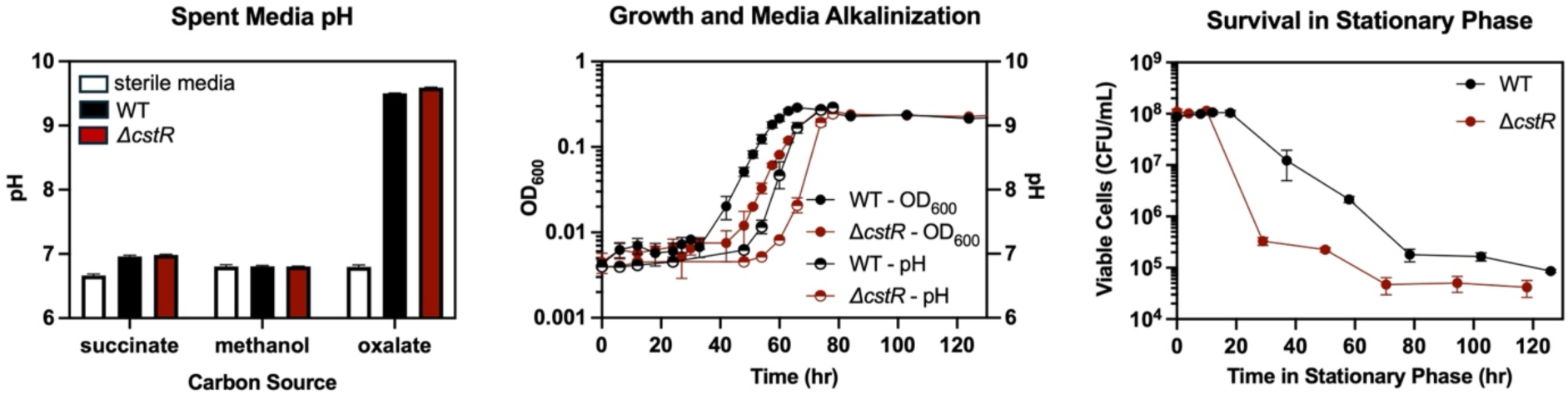
Growth and survival in oxalate media. A) Stationary phase media pH values were measured from MP media following cell growth of WT and Δ*cstR* strains upon either succinate, methanol, or oxalate as the sole carbon source. B) Percentage of population surviving in stationary phase of WT and Δ*cstR* strains in the same media as panel A, measured at 72 hr of growth. C) Oxalate-grown cells (that lost viability in stationary phase) were subcultured in oxalate media and demonstrate an offset in growth (filled circles) due to different starting inoculum resulting from differential die-off; supernatant pH (half-filled circles) was monitored over time to demonstrate alkalinization of media during active growth. D) Viability in spent, alkalinized media was measured once cultures reached stationary phase (determined by observing a measured peak/non-change in OD_600_). Plotted values represent a mean of four biological replicates, and error is represented by SEM.

**Figure S6.**
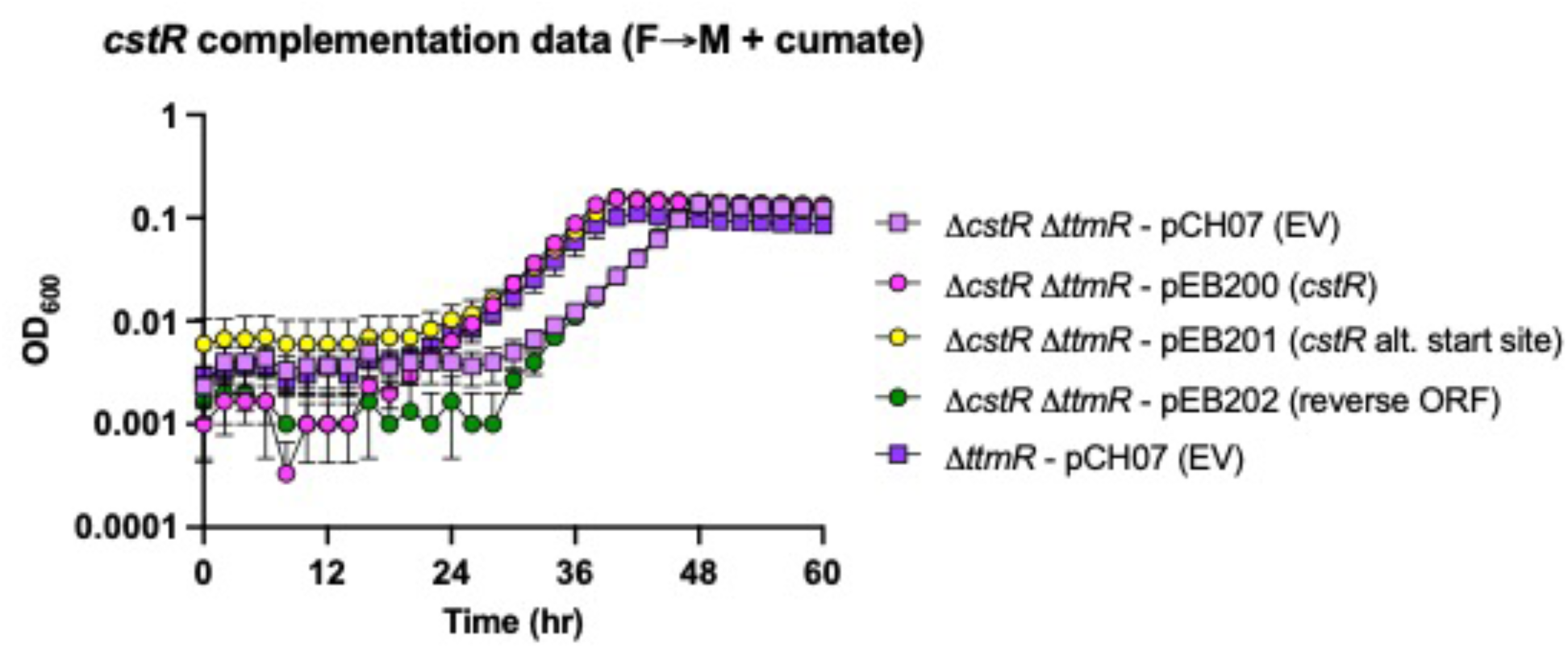
Complementation of CstR function is provided by the annotated CDS. Growth of strains lacking *ttmR* or *ttmR* and *cstR* and harboring a kanamycin-resistant expression plasmid under inducing conditions (containing cumate) was tracked during a formate-to-methanol transition. In addition to empty vector (EV) controls, plasmids containing a) annotated *cstR* (381 bp, pEB200), the extended *cstR* ORF (543 bp, pEB201), and a reverse ORF overlapping *cstR* (375 bp, pEB202); these ORFs are also visualized in **Figure S3**. Points represent the means of biological triplicates and error bars represent SEM.

**Table S1. Bacterial strains and plasmids.**

**Table S2. Population whole genome sequencing results.**

**Table S3. Differentially expressed genes in the Δ*cstR* background vs. WT *M.*** *extorquens*. The following list of genes were differentially regulated, as determined by a statistically significant change (FDR < 0.01) between *ΔcstR* and WT *M. extorquens* PA1 strains during growth with succinate as a sole carbon source or during the transition to methylotrophy (methanol as sole carbon source).

**Table S4. Gene enrichment among differentially expressed genes in the Δ*cstR* strain.**

